# Integrative analysis of genomic and transcriptomic alterations of *AGR2* and *AGR3* in cancer

**DOI:** 10.1101/2022.03.08.483441

**Authors:** Delphine Fessart, Ines Villamor, Eric Chevet, Frederic Delom, Jacques Robert

## Abstract

The *AGR2* and *AGR3* genes have been shown by numerous groups to be functionally associated to adenocarcinoma progression and metastasis. We explore in this paper the data available in databases concerning genomic and transcriptomic features concerning these two genes: the NCBI dbSNP database was used to explore the presence and roles of constitutional SNPs, and the NCI, CCLE and TCGA databases were used to explore somatic mutations and copy number variations (CNVs), as well as mRNA expression of these genes in human cancer cell lines and tumours. Relationships of *AGR2*/*3* expression with whole genome mRNA expression and cancer features (i.e. mutations and CNVs of oncogenes and tumour suppressor genes [TSG]) were established using CCLE and TCGA databases. In addition, the CCLE data concerning CRISPR gene extinction screens (Achilles project) were explored concerning these two genes and a panel of oncogenes and TSG. We observed that no functional polymorphism or recurrent mutation could be detected in *AGR2* or *AGR3*. The expression of these genes was positively correlated with the expression of epithelial genes and inversely correlated with that of mesenchymal genes. It was also significantly associated with several cancer features, such as *TP53* or *SMAD4* mutations, depending on the gene and the cancer type. The CRISPR screens revealed in addition the absence of cell fitness modification upon gene extinction, in contrast to oncogenes (cell fitness decrease) and TSG (cell fitness increase). Overall, these explorations revealed that AGR2 and AGR3 proteins appear as common non-genetic evolutionary factors in the process of human tumorigenesis.

## Introduction

Members of the protein disulphide isomerase (PDI) family, which are endoplasmic reticulum (ER)-resident enzymes interfering in the formation of disulphide bonds, cysteine-based redox reactions and quality control of proteins in the ER, play an essential role in ER homeostasis (proteostasis); in addition to their principal ER location, some of these enzymes are found in other localisations such as the extracellular milieu, in extracellular vesicles or the cytosol [1]. For instance, we have shown that PDIA2 is secreted into the lumen of the thyroid follicles by thyrocytes to control extracellular thyroglobulin folding and multimerisation [2, 3]. There is ample evidence supporting that PDI proteins are strongly associated with cancer either through their altered expression or through enhanced functions. Although they are among the most abundant cellular proteins, PDI expression is frequently up-regulated in cancers and associated with metastasis and invasiveness [1].

However, the functions of PDI proteins in the process of human oncogenesis remain to be understood. Among the most studied PDI in this respect are those belonging to the Anterior GRadient (AGR) family of proteins. The AGR family is composed of three proteins, namely AGR1 [gene *TXNDC12*], AGR2 and AGR3. Interestingly, AGR2, the prototypic member of the AGR family, is shown to play intracellular roles in the ER, contributing to proteostasis [4], but it remains unclear how this is related to oncogenesis. *AGR2* and *AGR3* genes are localised on chromosome 7, side by side (7p21.1), and their protein products are both overexpressed and their localisations deregulated in many types of adenocarcinomas [5–7]. We have shown that two non-canonical localisations: extracellular (eAGR2/3) [8, 9] [10] and cytosolic (cAGR2) [11] and exert pro-oncogenic gain-of-function to confer tumours specific and evolutive features (development, progression and aggressiveness). Moreover, the overexpression of AGR2 and AGR3 may be a prognosis factor for survival, which could be favourable or not favourable depending on the cancer type [7].

These observations raise the question of whether *AGR2* and *AGR3* could behave as ‘*cancer genes*’, i.e. as oncogenes and/or tumour suppressor genes (TSG). To bring some answers to this question, we have explored publicly available databases to search for relationships between genomic variations of *AGR2* and *AGR3* and cancer. In a first attempt, polymorphisms were sought in germline DNA using the dbSNP database; then, somatic tumour variations were sought in the TCGA tumour collection and in the CCLE cell line database, so as to elaborate a directory of potentially oncogenic mutations. In addition to the exploration of the sequence of these genes in constitutional and tumour DNA, we explored the expression pattern of both genes in tumour and cell lines of various tissue origins, and searched for relationships between *AGR2* and *AGR3* gene expression and several oncogenic determinants in various cancer types, tumours or cell lines, especially copy number variations (CNV) and point mutations (single nucleotide variations, SNV). We performed thus a comprehensive analysis of available data in order to better understand the role of AGR2 and AGR3 in cancer. All the analyses were conducted on the data available online in April 2021.

## Methods

### Databases

The dbSNP database was accessed from the NCBI database using the followings links:

For AGR2: https://www.ncbi.nlm.nih.gov/SNP/snp_ref.cgi?locusId=10551

For AGR3: https://www.ncbi.nlm.nih.gov/SNP/snp_ref.cgi?locusId=155465

We restricted our analysis to exomic variations. Synonymous variations were not studied. TCGA (The cancer Genome Atlas) was accessed through the cBioPortal for Cancer Genomics: https://www.cbioportal.org. Only data coming from the PanCancer Atlas were retrieved; they concern 32 different cancer types for a total of 10,945 tumours. Data concerning single nucleotide variations (SNV), copy number variations (CNV) and mRNA expression (RSEM, Batch normalized from Illumina HiSeq_RNASeqV2) were downloaded and converted into Excel sheets for analysis. We used the cancer type nomenclature of the TCGA (Supplemental Table 1). The CCLE (Cancer Cell Line Encyclopedia) was accessed through a friendly-user platform, https://discover.nci.nih.gov/cellminercdb/, established at NCI and gathering all publicly available data concerning cancer cell line molecular and pharmacologic properties [12, 13]. Rapid surveys of collections other than CCLE (namely GDSC, Genomics of Drug Sensitivity in Cancer, and CTRP, Cancer Therapeutics Response Portal) was performed in order to assess the accuracy of CCLE data. Most of the other analyses were conducted on the CCLE collection, which contained the highest number of cell lines, but all three collections are redundant and contain the same core cell lines, so that this restriction does not generate any bias.

**Table 1.**
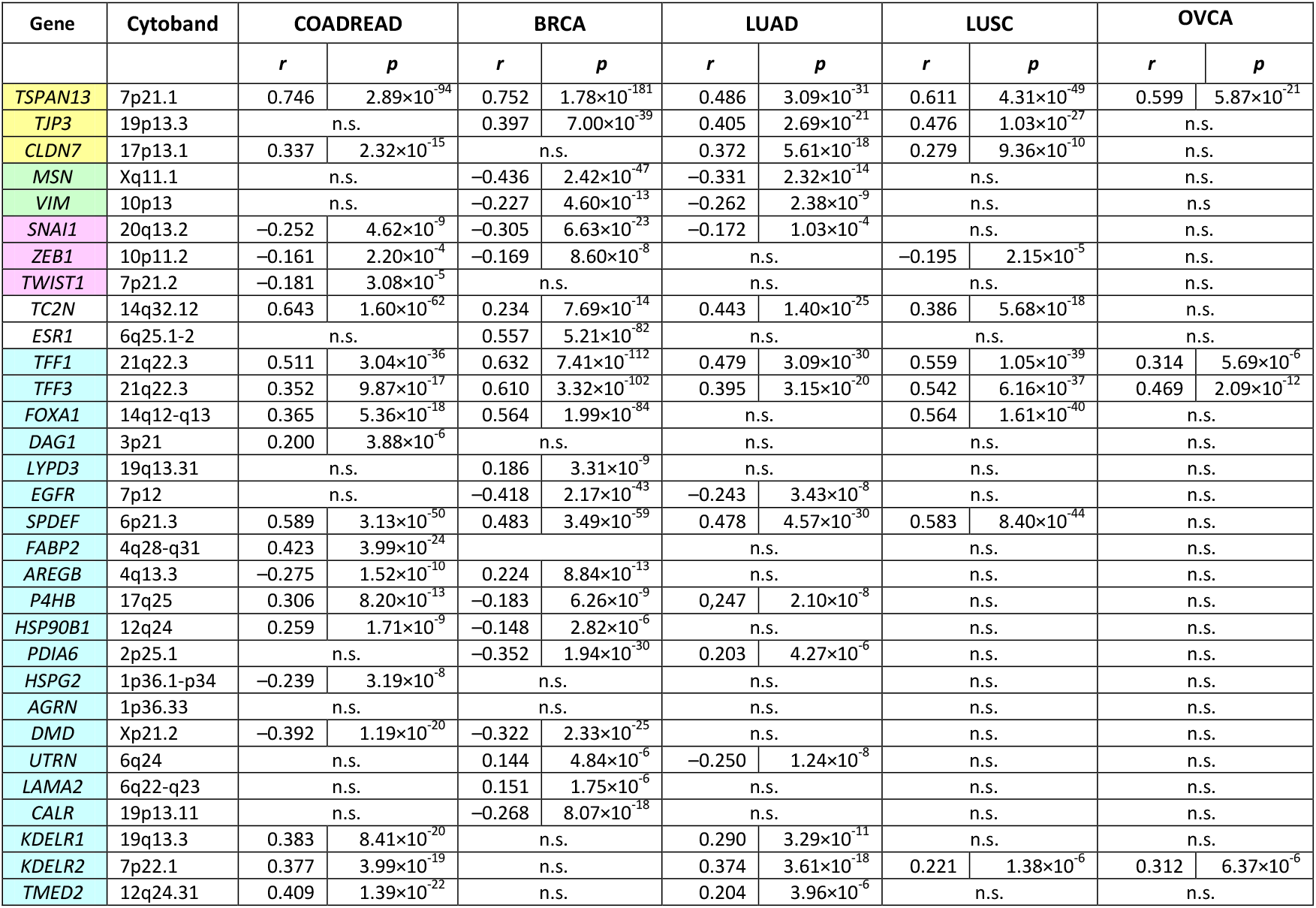
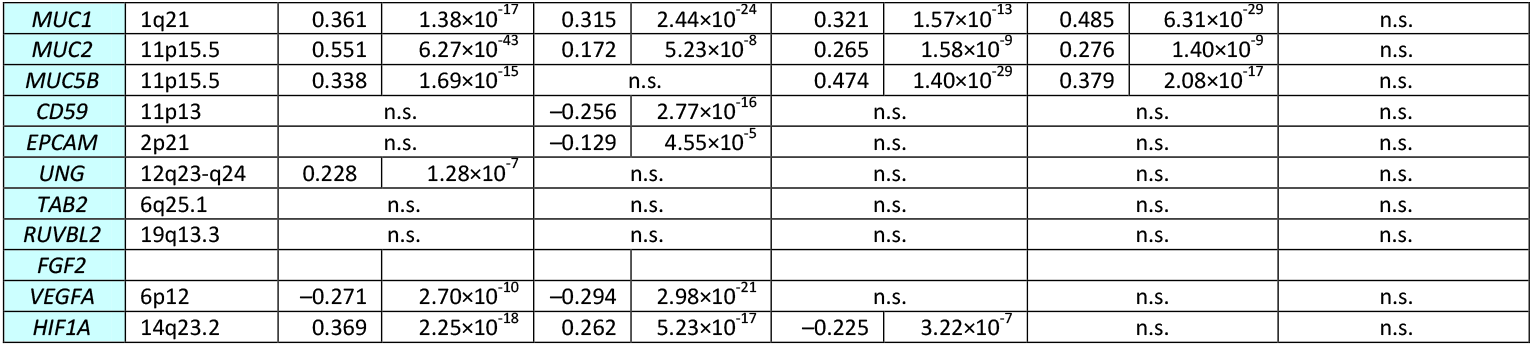
List of selected genes whose expression is correlated with that of *AGR2* in five major cancer types of TCGA. Gene selection was arbitrary; we have selected genes representative of epithelial features in yellow (*TJP3, TSPAN13, CLDN7*), of mesenchymal features in green (*MSN, VIM*), of EMT in pink (*SNAI1, ZEB1, TWIST1*) as well as *TC2N* and *ESR1*, which are already known to be associated with *AGR2* in colon and breast carcinomas, respectively. In addition, genes encoding proteins known to interact with AGR2 [14, 15] were studied (spotted in blue). Threshold for significance was set at 10^−8^ because of multiple testing, but we indicated *p* values down to 10^−4^ to indicate trends at the limit of significance. *r*: Pearson coefficient of correlation; *p*: degree of significance of the correlation.

### Statistics

We currently used common statistical tests for data comparisons, mainly Chi-2 and Student’s t test; all tests were two-sided and differences were considered that significance was obtained only at the 1% level. Large numbers of statistical tests were performed in several instances, and we took then multiple testing into account by applying the Bonferroni correction. For instance, as many as 12 × 20,000 *p* values were computed for gene association detection: in such cases, we considered only *p* < 5 × 10^−8^ as significant at the 1% level.

## Results

### AGR2 and AGR3 polymorphisms

In order to distinguish germline polymorphisms from potential mutations in tumour tissues, we first listed the *AGR2* and *AGR3* gene polymorphisms identified in the NCBI dbSNP database. In this database, 165 SNV or small insertion/deletion variations (indel) in the *AGR2* gene coding sequence are listed, affecting 115 of the 175 amino-acids of the protein. When indicated in the database, none of them has a minor allele frequency (MAF) higher than 0.0002, with the exception of rs6842 (N147N), a synonymous variation with a MAF of 0.3355. These variations were synonymous (41 cases), missense (112), nonsense (6), frameshift (7) or in frame (1).

Similarly, in the NCBI dbSNP database, 214 SNV or indels have been described in the *AGR3* gene coding sequence, affecting 131 of the 166 amino-acids of the protein. When indicated, none of them has a minor allele frequency (MAF) higher than 0.0006, with the exception of rs55900499 (D40D), a synonymous variation with a MAF of 0.0505. These variations were synonymous (48 cases), missense (151), nonsense (11), or frameshift (4).

### AGR2 and AGR3 somatic tumour variations in the TCGA

The TCGA database provides a unique comprehensive resource for exploring gene variations occurring in human tumours. Out of a total of 9,888 tumours originating from 32 tumour types (list and abbreviations in Supplemental Table 1), we identified 32 samples bearing an *AGR2* gene variation (mutation or polymorphism) (Figure 1A) and 35 bearing an *AGR3* gene variation (Figure 1B). A total of 30 different variations involving 26 codons in *AGR2*, and 31 mutations involving 26 codons in *AGR3*, were present in these samples. Three samples presented two variations in the *AGR2* sequence and two other samples in *AGR3* sequence (Figure 2). Only three samples showed variations in both *AGR2* and *AGR3* genes (Figure 2). Most variations were missense mutations; there were three nonsense mutations in *AGR2* and two in *AGR3*; two frameshift mutations in *AGR2* and one in *AGR3*; and two splice mutations in *AGR2* and one in *AGR3*. Some cancer types presented more mutations than other ones: skin cutaneous melanomas and endometrial carcinomas for *AGR2* (Supplemental Table 2A), and the same plus stomach and bladder carcinomas for *AGR3* (Supplemental Table 2B). Among the 30 *AGR2* variations found in the TCGA, seven were in the dbSNP list, 16 affected a codon where a SNP had been identified and seven concerned a codon not known as subject to polymorphic variation. Among the 31 *AGR3* variations found in TCGA, five were in the dbSNP list, 17 affected a codon where a SNP had been identified and nine concerned a codon not known as subject to a polymorphic variation.

**Table 2.** List of genes whose expression is highly positively or negatively correlated with that of *AGR2* in the whole set of cell lines of the CCLE collection. Gene selection was arbitrary; we have selected genes representative of epithelial features in yellow (*TJP3, TSPAN13, CLDN7*), of mesenchymal features in green (*MSN, VIM*), of EMT in pink (*SNAI1, ZEB1, TWIST1*) as well as *TC2N* and *ESR1*, which are already known to be associated with *AGR2* in colon and breast carcinomas, respectively. In addition, genes encoding proteins known to interact with AGR2 [14, 15] were studied (spotted in blue). Threshold for significance was set at 10^−8^ because of multiple testing, but we indicated *p* values down to 10^−4^ to indicate trends at the limit of significance. *r*: Pearson coefficient of correlation; *p*: degree of significance of the correlation.

| Gene |  | *AGR2* |  | *AGR3* |  |
| --- | --- | --- | --- | --- | --- |
|  |  | *r* | *p* | *r* | *p* |
| *TSPAN13* | 7p21.1 | 0.507 | $1.03 \times 10^{-68}$ | 0.337 | $6.95 \times 10^{-29}$ |
| *TJP3* | 19p13.3 | 0.736 | $4.94 \times 10^{-177}$ | 0.565 | $1.71 \times 10^{-88}$ |
| *CLDN7* | 17p13.1 | 0.719 | $2.05 \times 10^{-165}$ | 0.480 | $6.61 \times 10^{-61}$ |
| *MSN* | Xq11.1 | -0.521 | $3.39 \times 10^{-73}$ | -0.494 | $7.62 \times 10^{-65}$ |
| *VIM* | 10p13 | -0.581 | $1.19 \times 10^{-94}$ | -0.448 | $2.67 \times 10^{-52}$ |
| *SNAI1* | 20q13.2 | -0.181 | $4.89 \times 10^{-9}$ | -0.143 | $4.05 \times 10^{-6}$ |
| *ZEB1* | 10p11.2 | -0.592 | $4.20 \times 10^{-99}$ | -0.428 | $2.05 \times 10^{-47}$ |
| *TWIST1* | 7p21.2 | -0.285 | $8.20 \times 10^{-21}$ | -0.187 | $1.32 \times 10^{-9}$ |
| *TC2N* | 14q32.12 | 0.727 | $4.95 \times 10^{-171}$ | 0.556 | $3.53 \times 10^{-85}$ |
| *ESR1* | 6q25.1-2 | 0.196 | $2.08 \times 10^{-10}$ | 0.170 | $3.85 \times 10^{-8}$ |
| *TFF1* | 21q22.3 | 0.701 | $9.50 \times 10^{-154}$ | 0.592 | $5.15 \times 10^{-99}$ |
| *TFF3* | 21q22.3 | 0.460 | $2.89 \times 10^{-55}$ | 0.542 | $4.37 \times 10^{-80}$ |
| *FOXA1* | 14q12-q13 | 0.669 | $2.79 \times 10^{-135}$ | 0.432 | $2.67 \times 10^{-48}$ |
| *DAG1* | 3p21 | 0.249 | $4.02 \times 10^{-16}$ | 0.123 | $6.79 \times 10^{-5}$ |
| *LYPD3* | 19q13.31 | 0.428 | $1.99 \times 10^{-47}$ | 0.166 | $7.04 \times 10^{-8}$ |
| *EGFR* | 7p12 | 0.270 | $9.01 \times 10^{-19}$ | n.s. | |
| *SPDEF* | 6p21.3 | 0.471 | $2.69 \times 10^{-58}$ | 0.340 | $1.77 \times 10^{-29}$ |
| *FABP2* | 4q28-q31 | n.s. |  | n.s. |  |
| *AREGB* | 4q13.3 | 0.430 | $5.47 \times 10^{-48}$ | 0.269 | $1.26 \times 10^{-18}$ |
| *P4HB* | 17q25 | n.s. |  | n.s. |  |
| *HSP90B1* | 12q24 | n.s. |  | n.s. |  |
| *PDIA6* | 2p25.1 | n.s. |  | n.s. |  |
| *HSPG2* | 1p36.1-p34 | n.s. |  | n.s. |  |
| *AGRN* | 1p36.33 | 0.328 | $1.92 \times 10^{-27}$ | n.s. | |
| *DMD* | Xp21.2 | -0.174 | $1.66 \times 10^{-8}$ | n.s. | |
| *UTRN* | 6q24 | n.s. | | -0.147 | $1.9 \times 10^{-6}$ |
| *LAMA2* | 6q22-q23 | n.s. |  | n.s. |  |
| *CALR* | 19p13.11 | n.s. | | -0.134 | $1.42 \times 10^{-5}$ |
| *KDELR1* | 19q13.3 | 0.193 | $4.07 \times 10^{-10}$ | n.s. | |
| *KDELR2* | 7p22.1 | 0.191 | $6.32 \times 10^{-10}$ | n.s. | |
| *TMED2* | 12q24.31 | 0.141 | $5.28 \times 10^{-6}$ | n.s. | |
| *MUC1* | 1q21 | 0.505 | $3.87 \times 10^{-68}$ | 0.382 | $2.01 \times 10^{-37}$ |
| *MUC2* | 11p15.5 | 0.393 | $1.16 \times 10^{-39}$ | 0.452 | $3.21 \times 10^{-53}$ |
| *MUC5B* | 11p15.5 | 0.387 | $2.73 \times 10^{-38}$ | 0.296 | $2.25 \times 10^{-22}$ |
| *MUC5AC* | 11p15.5 | 0.401 | $2.69 \times 10^{-41}$ | 0.266 | $2.68 \times 10^{-18}$ |
| *CD59* | 11p13 | n.s. | | -0.124 | $6.28 \times 10^{-5}$ |
| *EPCAM* | 2p21 | 0.641 | $3.46 \times 10^{-121}$ | 0.427 | $2.93 \times 10^{-47}$ |
| *UNG* | 12q23-q24 | n.s. | | 0.144 | $3.42 \times 10^{-6}$ |
| *TAB2* | 6q25.1 | n.s. |  | n.s. |  |
| *RUVBL2* | 19q13.3 | n.s. |  | n.s. |  |
| *FGF2* | 4q26 | -0.348 | $6.07 \times 10^{-31}$ | -0.305 | $8.17 \times 10^{-24}$ |
| *VEGFA* | 6p12 | n.s. |  | n.s. |  |
| *HIF1A* | 14q23.2 | n.s. | | -0.132 | $2.07 \times 10^{-5}$ |

**Figure 1.**
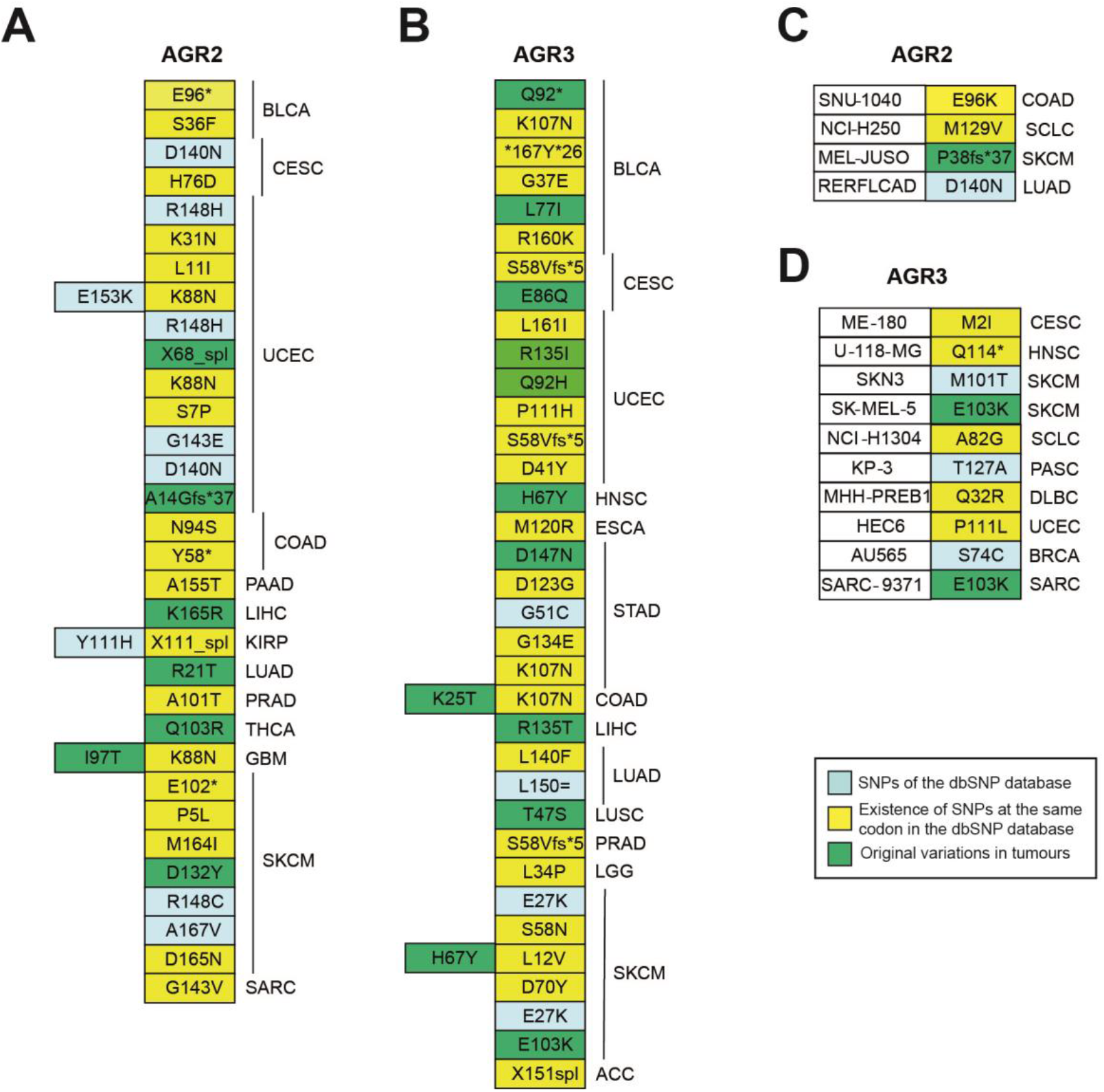
Point mutations of *AGR2* and *AGR3* present in databases. **A-B**, Point mutations of *AGR2* (A) and *AGR3* (B) genes in 10,376 tumour samples of the TCGA. The standard Mutation Nomenclature in Molecular Diagnostics can be found at https://www.hgvs.org/mutnomen/recs-prot.html. **C-D**, Point mutations in *AGR2* (C) and *AGR3* (D) genes in 1036 cell lines of the CCLE and GDSC collections

**Figure 2.**
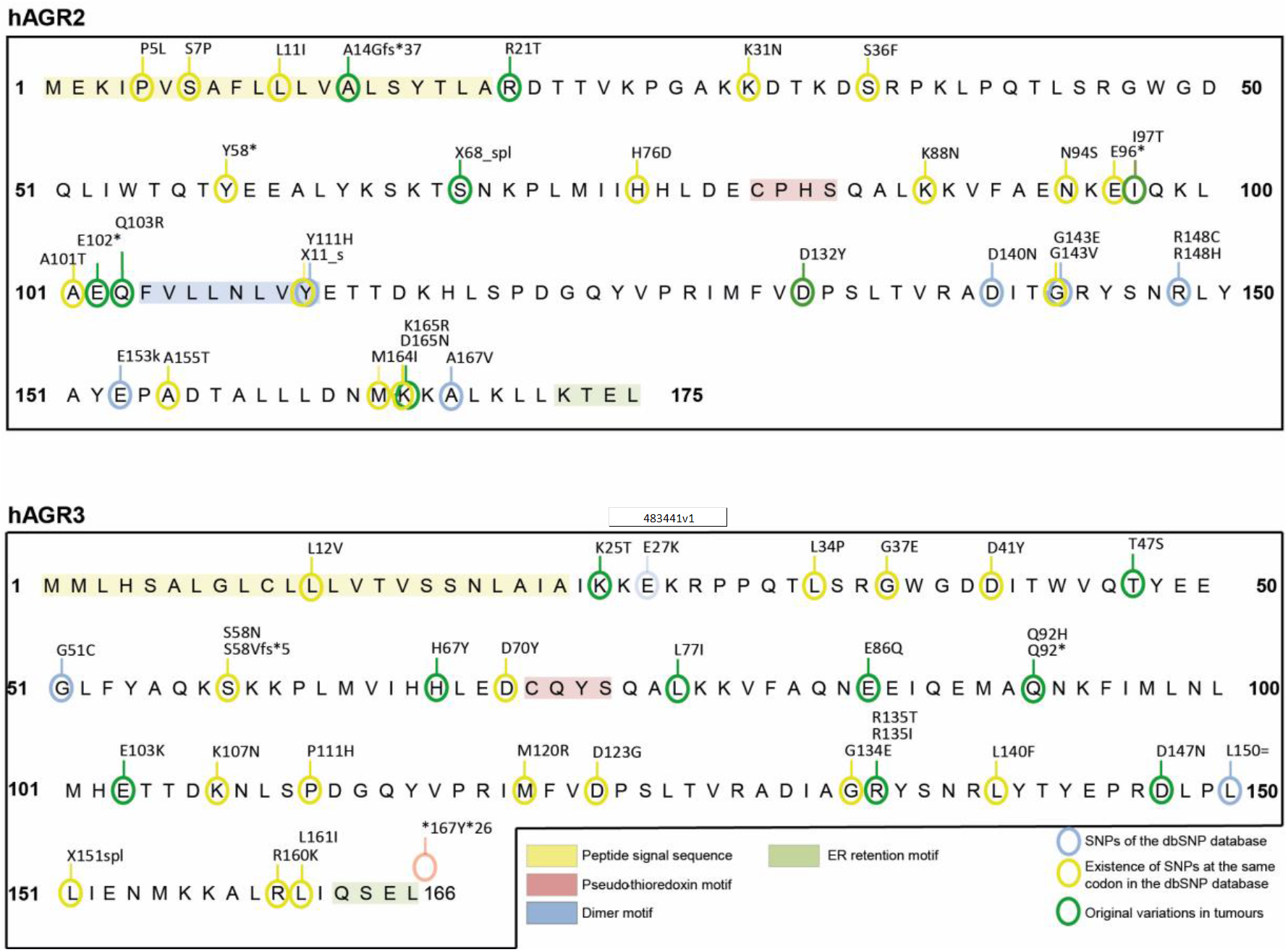
Localisation of constitutional SNPs and tumour somatic SNVs in the sequence of AGR2 and AGR3 proteins. The functional domains of the proteins are indicated.

Concerning copy number variations, there were in TCGA 146 samples with *AGR2* gene amplifications and 14 with *AGR2* homozygous deletion (Supplemental Table 2A); and 145 samples with *AGR3* gene amplification and 15 with *AGR3* homozygous deletion (Supplemental Table 2B). Most of samples were amplified on both genes, nine samples presenting *AGR2* amplification only and seven *AGR3* amplification only. Similarly, only one sample had a homozygous deletion of only one of the two genes, *AGR3*.

### AGR2 and AGR3 somatic tumour variations in cell line collections

In the collections of cell lines of GDSC (Genomics of Drug Sensitivity in Cancer) and CCLE, four tumour cell lines bear a variation in *AGR2* coding sequence, among which three are common to the two databases. One is listed in the NCBI dbSNP database, two occur at a codon where other SNPs are listed in the database and one is original (P38fs*37) (Figure 1C).

In the collections of cell lines of GDSC and CCLE, 10 tumour cell lines bear a variation in *AGR3* coding sequence, among which five are common to the two databases. Three are listed in the NCBI dbSNP database, five occur at a codon where other SNPs are listed in the database and one is original (E103K) and present in two cell lines SK-MEL-5 (human melanoma cell line) and SARC-9371 (human osteosarcoma cell line) (Figure 1D).

Figure 2 presents the localisation of constitutional SNPs and tumour somatic SNVs extracted from the CCLE and TCGA databases, in the sequence of AGR2 and AGR3 proteins. With the exception of some known SNPs, none of them is present in the functional domains of the proteins.

### AGR2 and AGR3 expression in TCGA and CCLE databases

Thanks to the cBioPortal facilities for TCGA and the CellMinerCDB portal for CCLE and GDSC, it was possible: (**i**) to compare the levels of *AGR2* and *AGR3* expressions in various tumour types; (**ii**) to identify *AGR2*/*3* expression variations in samples with SNV or CNV of these genes; (**iii**) to look for associations between *AGR2*/*3* expression and that of other genes in selected tumour types; (**iv**) to identify associations between *AGR2*/*3* and potentially oncogenic molecular features involving the whole exome, namely SNV and CNV. Since *AGR2* and *AGR3* expressions were highly correlated (Figure 3) in all the TCGA tumour types studied as well as in the CCLE collection, we focused our interest on *AGR2* and simply indicated original features concerning *AGR3*.

**Figure 3.**
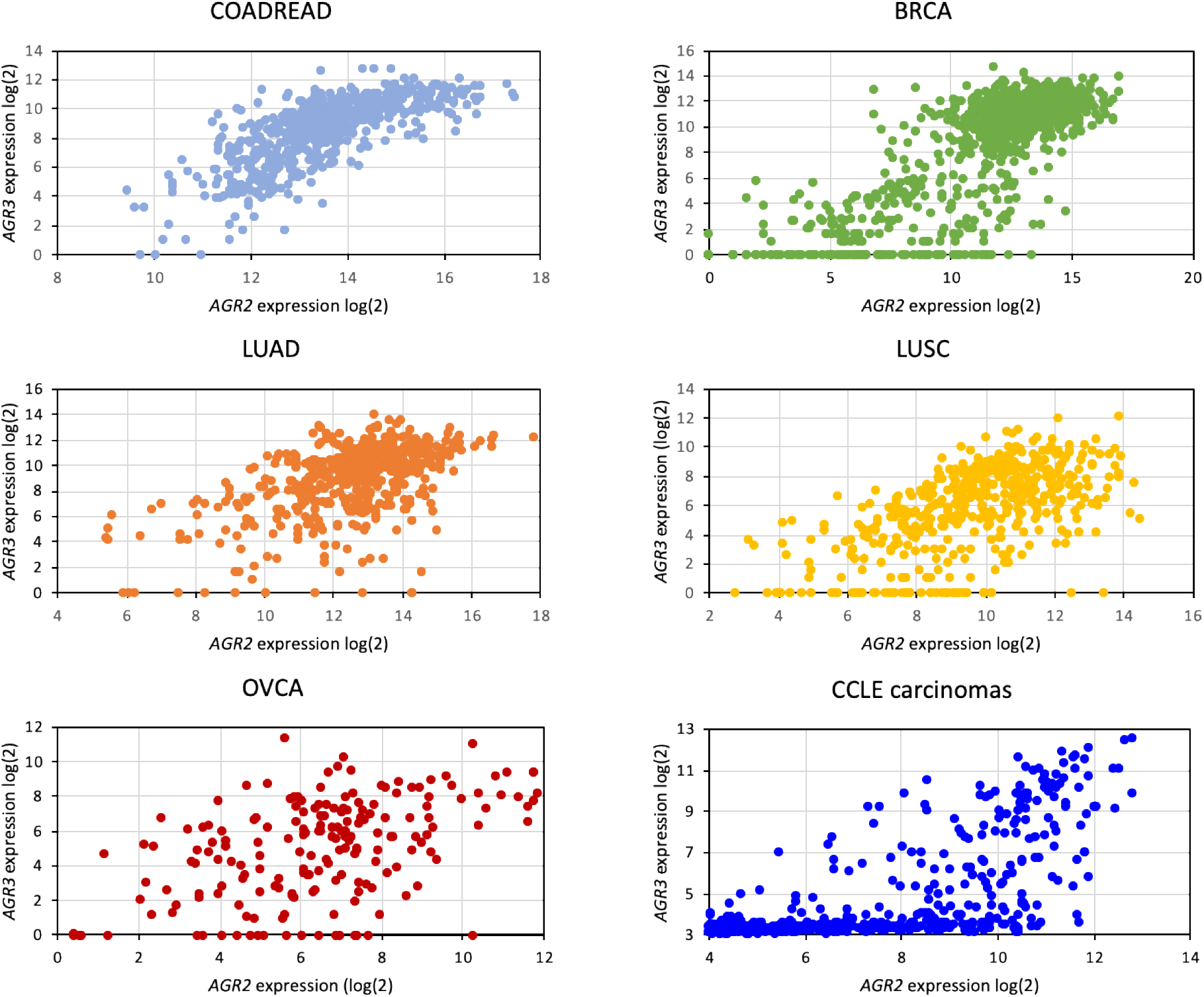
Correlation between *AGR2* and *AGR3* mRNA expressions in five TCGA tumour types (COADREAD, BRCA, LUAD, LUSC, OVCA) and in the 634 carcinoma cell lines from the CCLE.

#### (i) Expression levels

Among the 32 cancer types that are available in the PanCancer Atlas project of TCGA, only part of them display a consistent expression of *AGR2* and *AGR3*. Non-epithelial cancers do not express this gene, and carcinomas from liver and kidney express these genes in a small part of the samples only, not always distinguishable from background noise; as a consequence, we concentrated our analysis on BLAD, CESC, UCEC, HNSC, STAD, ESCA, LUAD, LUSC, COADREAD, PAAD, PRAD, BRCA, and OVCA (Figure 4A). In all cancer types, *AGR3* was expressed at a lower level than *AGR2*, and often not evaluable in samples from three cancer types: BLCA, HNSC, and OVCA. The expression levels of the two genes were highly correlated in each cancer type. As a general feature, squamous cell carcinomas expressed *AGR2* and *AGR3* at a much lower level than adenocarcinomas (compare, for instance, LUAD with LUSC, ESCA with HNSC, CESC with UCEC).

**Figure 4.**
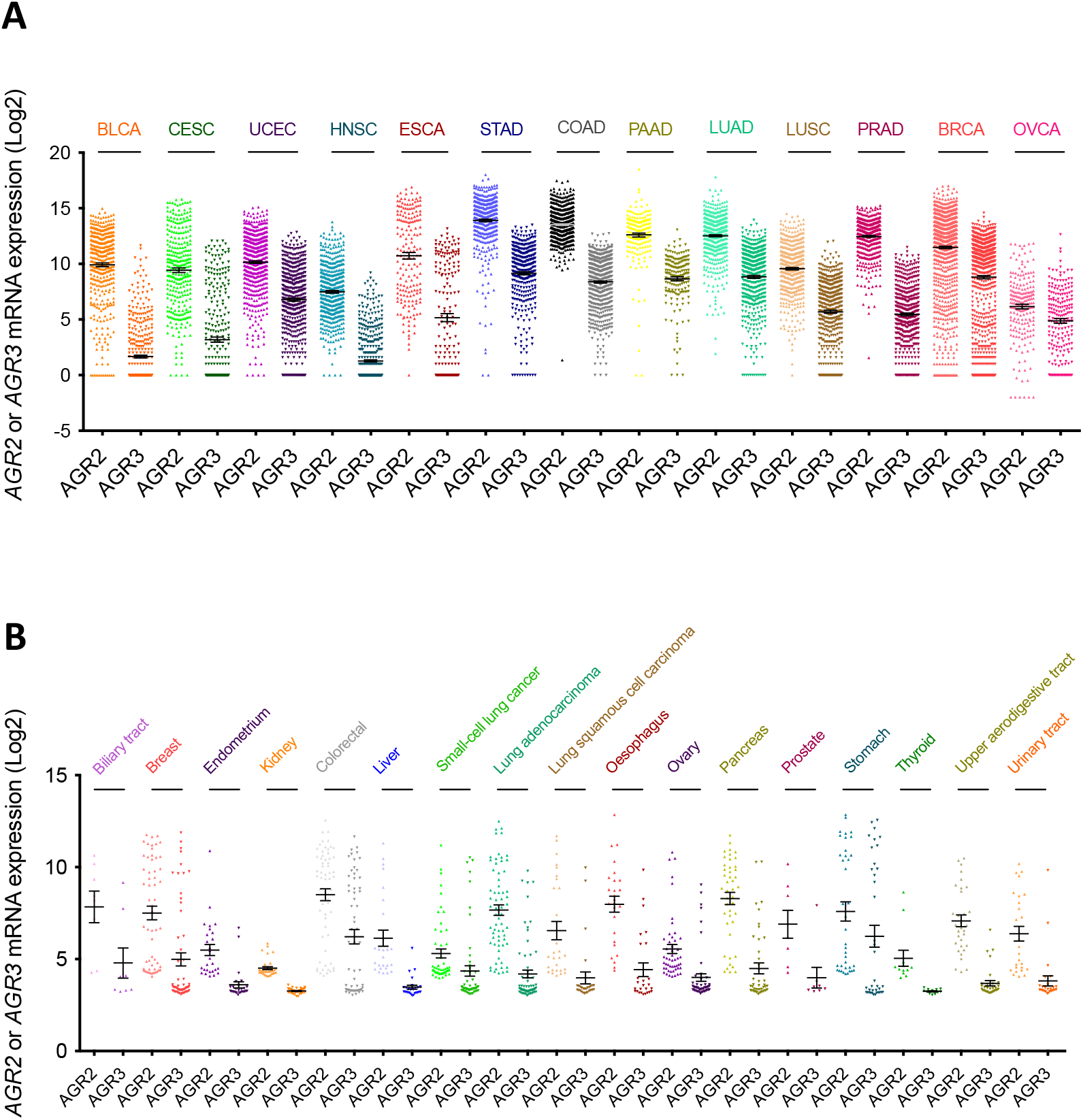
mRNA expression levels of *AGR2* and *AGR3* extracted from databases. **A**, expression in 13 major cancer types from the TCGA database. **B**, expression in 17 cancer cell types from the CCLE database.

In the CCLE collection, the levels of expression of *AGR2* and *AGR3* also vary considerably across cancer types. As a general feature, cancer cells derived from mesenchymal tissues express these genes to low levels, barely higher than background noise, whereas cancer cells derived from epithelial tissues have consistent expression levels. As a consequence, we excluded from further analyses cancer cells from autonomic ganglia (neuroblastoma), bone (osteosarcoma and Ewing’s sarcoma), central nervous system (glioma), haematopoietic and lymphoid tissue, pleura, skin (malignant melanoma), and soft tissue sarcomas. Figure 4B presents the levels of expression of *AGR2* and *AGR3* in all other cancer cell line types. Cell lines derived from digestive tract cancers (with the exception of liver) had the highest expression levels, while cell lines derived from cancers of kidney, endometrium, ovary and thyroid carcinomas had the lowest expression levels.

#### (ii) *AGR2/3* expression variation

In the TCGA, the expression of *AGR2* and *AGR3* in genomic variants of these genes was not markedly different from that mentioned for the unaltered samples. Concerning CNV, looking for associations between *AGR2* or *AGR3* expression and copy number in five major tumour types (COADREAD, BRCA, LUAD, LUSC, and OVCA), revealed no significant correlations between these two parameters (data not shown). In addition, when considering SNV, nonsense or frameshift mutations in gene sequence were not associated to loss of gene expression. Another way of analysing relationships between CNV and expression was to consider chromosome 7p losses in these cancer types; there were only three shallow 7p deletions in COADREAD out of 492 samples, not allowing comparisons, but in BRCA (66 samples with 7p loss out of 850 samples) there was significantly lower *AGR2* and *AGR3* expressions when chromosome 7p was lost (*p* = 4.72 × 10^−13^ and 2.7 × 10^−9^ respectively); in LUAD and OVCA, barely significant lower expression values were noticed, and no significant results were obtained in LUSC (Figure 5).

**Figure 5.**
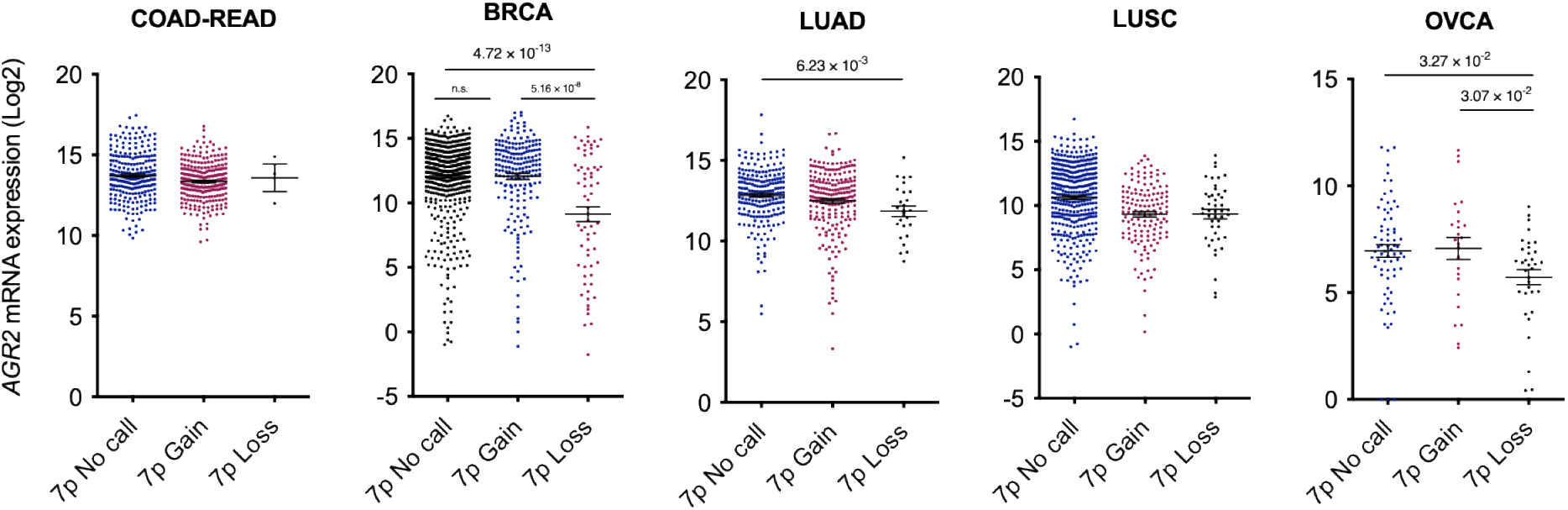
Association between chromosome 7p gains and losses and *AGR2* mRNA levels in five cancer types: BRCA, COAD-READ, OVCA, LUAD and LUSC.

In the CCLE collection, there was no clear association between the presence of *AGR2*/*3* sequence variations in cell lines and the expression of these genes. In the MEL-JUSO melanoma cell line, the frameshift P38fs*37 AGR2 variation is accompanied by the lowest *AGR2* mRNA expression in melanoma cell lines, but only in the GDSC database. No other peculiarities could be discerned. In contrast, there was a significant correlation between *AGR2* expression and gene copy number (*p* = 1.83 × 10^−9^) when the whole set of cell lines was taken into consideration; however, this significance was lost when individual cancer types was studied in this respect, due to the relatively low number of cell lines in each cancer type.

#### (iii) Associations with cancer genes

In the TCGA, we also identified the genes that were co-expressed with *AGR2* or *AGR3* in five major tumour types (COADREAD, BRCA, LUAD, LUSC, and OVCA). Each of them had a specific set of genes positively and negatively associated with that of *AGR2*/*3*. In Table 1, we present the significance level of the correlations between *AGR2* expression and that of selected representative genes. As a general feature, the expression of epithelial genes (e.g. *TJP3, TSPAN13, CLDN7, EPCAM*) was positively correlated with the expression of *AGR2*/*3* and the expression of mesenchymal genes (e.g. *VIM, MSN*) was negatively correlated, with specific correlations according to cancer type. The expression of the genes encoding the transcription factors involved in EMT (*SNAI, ZEB* and *TWIST* families) were often negatively correlated with *AGR2* expression, but this generally remains slightly below the level of significance we have chosen for 1% risk. It was remarkable that *ESR1* (oestrogen receptor) was highly significantly associated with *AGR2* in BRCA, but not in other malignancies. Similarly, *FOXA1* and *AGR2* expressions were correlated in BRCA, COADREAD and LUSC, but not in LUAD or OVCA. The expression of genes encoding mucins (*MUC1, MUC2, MUC5A*) or involved in mucosa protection (*TFF1* and *TFF3*) were positively correlated with *AGR2* expression in most tumour types. In addition, genes encoding proteins known to interact with AGR2 [14, 15] were studied. There was a clear specificity in their co-expression pattern with *AGR2*: some genes were co-expressed in colon adenocarcinoma, other in breast adenocarcinoma, etc. It should be mentioned that *EGFR, CD59* and *VEGFA* gene expressions were in contrast negatively correlated with *AGR2* expression in breast adenocarcinomas.

In the CCLE collection as in TCGA, the genes significantly positively co-expressed with *AGR2* and *AGR3* in the whole set of 1036 cell lines of the CCLE were mostly epithelial genes, according to the list established by Kohn et al. [16]. Conversely, the expression of mesenchymal genes was inversely correlated with *AGR2* and *AGR3* gene expressions (Table 2). This is not surprising, in view of the fact that these genes were expressed to a much higher level in epithelial tissue-derived cell lines than in mesenchymal tissue-derived ones. However, when cancer types were studied independently (namely breast, colorectal, lung and ovarian adenocarcinomas), the same positive correlation between *AGR2* and *AGR3* expressions and those of epithelial genes was maintained, as well as the negative correlation between *AGR2* and *AGR3* expressions and those of mesenchymal genes (data not shown). In addition to epithelial/mesenchymal genes, some interesting associations could be identified: *AGR2* and *AGR3* mRNA levels are positively associated with high significance with *FOXA1* expression, *TFF1*/*2*/*3*, and *ESR1*. It is interesting to note that the expressions of genes encoding the transcription factors of EMT are negatively correlated with those encoding *AGR2* and *AGR3*: *ZEB1*/*2* with a very high significance, *TWIST1*/*2* and *SNAI1*/*2* with lower *p* values, but still highly significant. The genes encoding AGR2 protein interactants were positively co-expressed with *AGR2* for some of them such as *KDELR, TMED2, DAG1, LYPD3, MUC1*/*2*/*5AC*/*5B*) but negatively correlated for others such as *DMD* or *FGF2*. For AGR3 interactants, a distinct pattern was observed, with positive co-expressions with *DAG1, LYPD3, MUC1*/*2*/*5AC*/*5B* or *UNG*, and negative correlations with *UTRN, CALR, CD59, FGF2* or *HIF1A*.

#### (iv) Association with oncogenic features

We eventually wanted to know whether some oncogenic alterations in various pathways were associated to *AGR2* and *AGR3* expressions. Indeed, the oncogenic status of these genes is not clear and the possible association with established oncogenic features could shed some light upon this status. In this respect, we have selected in the TCGA the five tumour types (COADREAD, BRCA, LUAD, LUSC, and OVCA) and the set of genes which are the most commonly mutated in these malignancies (*KRAS, APC, TP53, SMAD4, BRAF* and *PIK3CA* for COADREAD; *TP53, PIK3CA, BRCA1*/*2* and *PTEN* for BRCA; *KRAS* and *TP53* for LUAD and LUSC; *TP53, BRCA1*/*2* and *RB1* for OVCA).

Concerning COADREAD (Figure 6A), it appeared that the presence of a *KRAS* mutation in a tumour was not associated with *AGR2* expression, whereas the presence of *APC* or *TP53* mutation was negatively associated with *AGR2* expression, and the presence of *SMAD4, BRAF* or *PIK3CA* mutation are positively associated with *AGR2* expression. It was the same for *MTOR, MLH1* and *MSH2* mutations (data not shown). Very similar associations were found between *AGR3* expression and oncogenic mutations in COADREAD, the only differences being the exact level of significance (data not shown).

**Figure 6.**
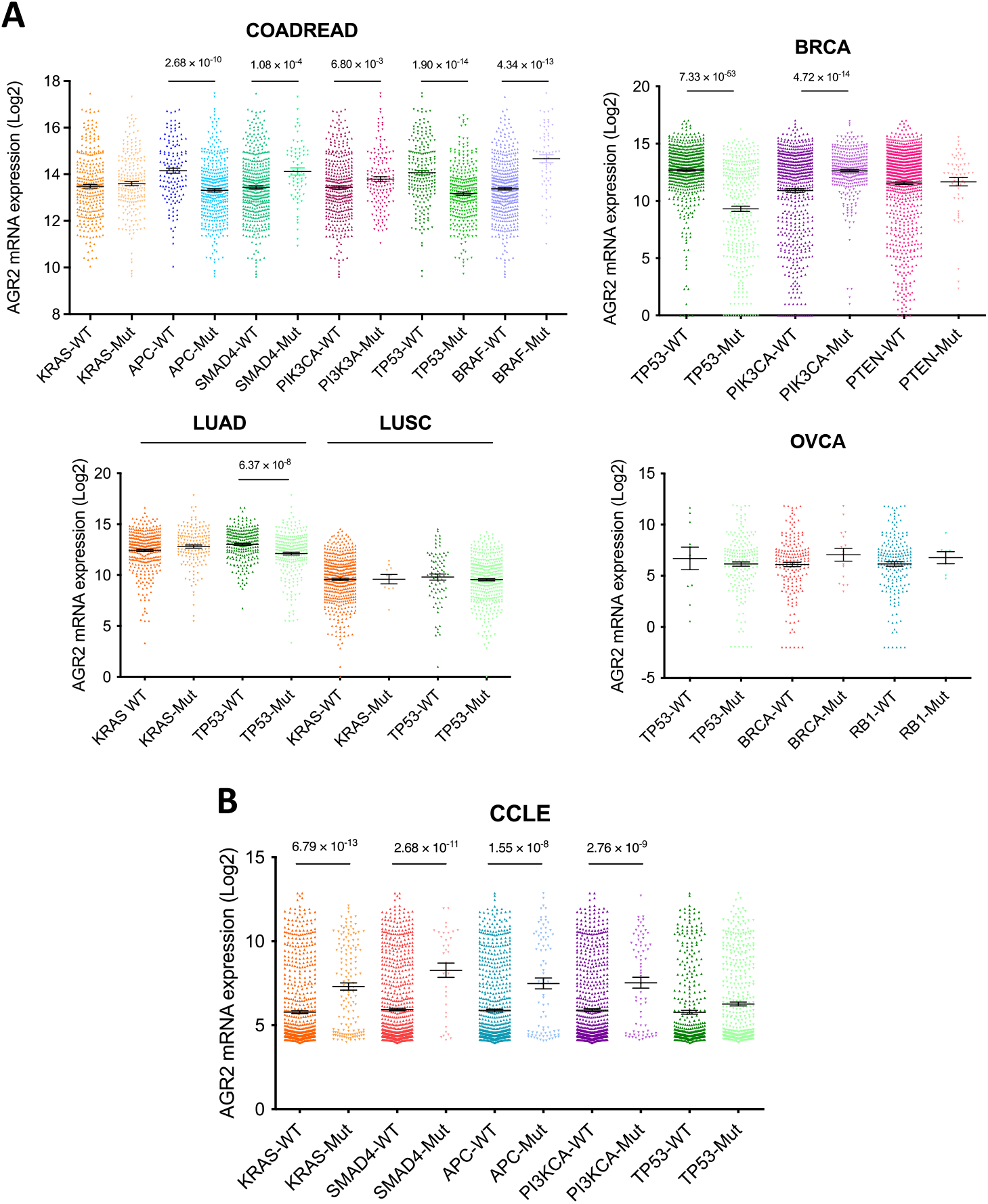
*AGR2* gene expression levels are associated to oncogene and TSG mutation status in different cancer types from the TCGA database and the CCLE database. **A**, Box plots displaying *AGR2* expression levels in COADREAD, BRCA, LUAD, LUSC and OVCA tumours with a mutation in genes *KRAS, APC, SMAD4, PIK3CA, BRAF, TP53, PTEN, BRCA*, and *RB1. p*-values were assessed using Student’s *t*-test. **B**, Box plots displaying *AGR2* expression levels in cancer cell lines with a mutation in genes *KRAS, SMAD4, APC, PIK3CA*, and *TP53. p*-values were assessed using Student’s *t*-tests.

Concerning BRCA (Figure 6A), the same was observed for *TP53* and *PIK3CA*: negative association between *AGR2* expression and *TP53* mutations, positive association for *PIK3CA*; no significant association was found between *PTEN* or *BRCA1*/*2* mutations and *AGR2* expression. Concerning lung tumours (Figure 6A), there was no significant association between *KRAS* mutations and *AGR2* expression, while there was, as in COADREAD and BRCA, a negative association between *TP53* mutation and *AGR2* expression in LUAD samples (but this was not the case in LUSC samples). No association between *AGR2* expression and oncogenic mutations were noticed in OVCA (Figure 6A). There again, similar associations were found between *AGR3* expression and oncogenic mutations in these cancer types (data not shown).

In the CCLE taken as a whole, an increase in *AGR2* and *AGR3* expressions was systematically associated with several oncogenic mutations (Supplemental Table 3). As an illustration, we present in figure 6B the significative associations existing between the expressions of *AGR2* and the presence of representative oncogene and TSG mutations, namely those occurring in *KRAS, SMAD4, APC*, and *PIK3CA*. However, this significance was lost when individual cancer types were studied in this respect, due to the relatively low number of cell lines in each cancer type.

Looking further into the associations that could be found between *AGR2* or *AGR3* gene expression and oncogenic features, we analysed also the relationships between *AGR2* and *AGR3* expressions and the copy number variation (CNV) of a set of oncogenes and TSG that are activated in cancers by copy gains (including amplifications) and losses (including deletions) respectively.

In the COADREAD samples of TCGA, a significant correlation is obvious between *AGR2* expression and *FOXA1* expression, in relation to the correlation observed between *AGR2* gene expression and *FOXA1* copy number. Also, a significant change in *AGR2* gene expression accompanied several CNV features known to drive colorectal cancers, especially those involved in cell cycle control (*TP53, FBXW7, RB1, CDC27, AURKA*), in WNT signalling (*APC, WNT4, FZD3, AJUBA*) and others (*SMAD4, SMAD2*). Figure 7A presents a selection of representative associations and Supplemental Table 4A a list of significant associations (down to p < 10^−4^) between oncogene or TSG gene copy numbers and *AGR2* expression in COADREAD. Some oncogenes and TSG of this list are not known to be frequently altered in colorectal cancers; it should be noticed that they belong to 14q or 18q chromosome arms, which respectively harbour *FOXA1* and *SMAD2*/*4*, suggesting that this correlation might be in fact related to the same event of gain or loss of a whole chromosome arm and has no functional meaning. Very similar results were obtained with *AGR3* expression (data not shown), slight differences occurring for the genes that were just below or just above the limit of significance chosen (10^−4^).

**Figure 7.**
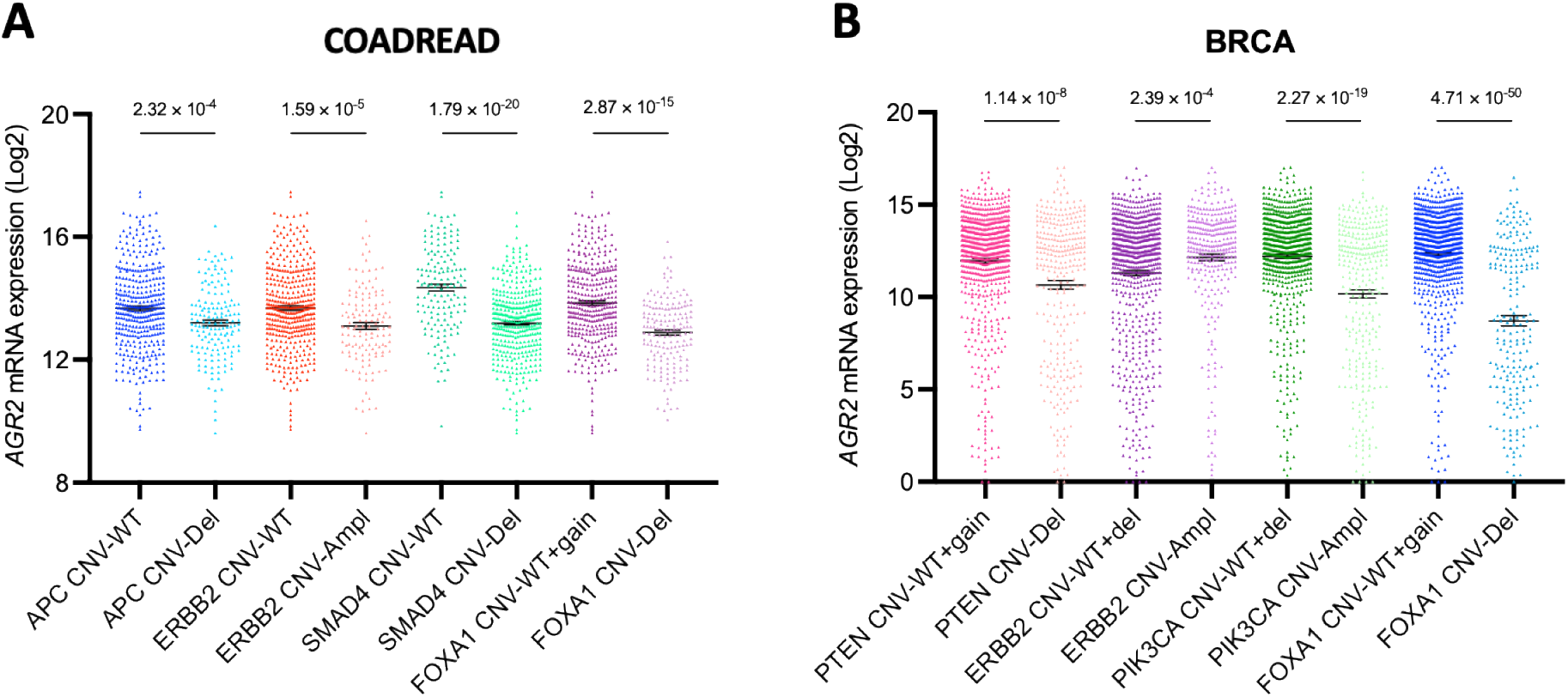
*AGR2* gene expression levels are associated to oncogene CNV in COADREAD and BRCA samples from the TCGA database. Only some examples are given, concerning principally genes known as oncogenic drivers in these cancer types; see Supplemental Table 4A for more details.

In the BRCA samples of TCGA, we also noticed a significant relationship between *AGR2* expression and *FOXA1* gene copy number, as well as several cancer genes copy numbers localised at 14q such as *NFKBIA, SAV1, CHD8* or *AJUBA*, which are not known as driver oncogenes or TSG in breast cancer (Figure 7B and Supplemental Table 4A). A highly significant association of *AGR2* expression was seen with *APC, JUN, CCNE1, ERBB2, MDM2* or *RAD21* copy numbers, which may have more functional implications. In contrast, copy numbers of *RB1* or *TP53* were not associated with *AGR2* expression, showing that the relationship between *AGR2* expression and oncogenic features in breast cancer is certainly complex and requires more in-depth analysis. Similar results were obtained with *AGR3* gene expression (data not shown), the differences between the two genes appearing to be marginal. No significant relationship between *AGR2* or *AGR3* expression and oncogene or TSG CNV was observed in LUAD, LUSC and OVCA (data not shown).

In the CCLE collection, CNV were not classified as gains or losses but copy numbers were given; we observed positive correlations between *AGR2* expression and gene copy numbers of several oncogenes such as *FOXA1, ERBB2, CCND1*, and *MYC*, whereas a negative correlation was found between *AGR2* expression and copy numbers of several TSG such as *SMAD4* (Supplemental Table 4B). However, this general trend was not constant over the whole set of oncogenes and GST. Similar results were obtained for *AGR3* with a lower number of cancer genes whose CNVs were associated with *AGR3* than with *AGR2* expression. In both cases, there was an overrepresentation of genes located on 14q and 19p chromosome arms, which may indicate that the association concerns a whole chromosome arm and not specific cancer genes. There again, this significance was lost when individual cancer types of the CCLE were studied, due to the relatively low number of cell lines in each cancer type.

##### Pattern of AGR2 extinction in the CCLE as studied by CRISPR screens

The Broad Institute has been set up CRISPR screens to study vulnerability targets through gene extinction screens in 769 cell lines of the CCLE collection [17]. It integrates data obtained by knocking-out each gene of the genome to analyse its consequences on cell viability and proliferation (regrouped as ‘cell fitness’). A friendly-user access has been made available by NCI on the CellMinerCDB site. The pattern of *AGR2* and *AGR3* gene extinction over cell lines can therefore be extracted and compared to the extinction pattern of other genes. The pattern of cell fitness alterations associated with *AGR2* and *AGR3* extinction are highly correlated (r = 0.331, *p* = 4.58 × 10^−21^) and did not reveal any preferential vulnerability towards a given cancer type represented in the cell line panels of the CCLE. No preferential effect was seen in epithelial vs. mesenchymal cell lines or in adenocarcinoma vs. squamous cell carcinomas, as was the case for expression data. The mean values of cell fitness alteration over 769 cell lines after *AGR2* and *AGR3* extinction are 1.003 ± 0.088 and 1.090 ± 0.075, whereas the same parameter is largely lower than 1 when oncogenes are knocked out (e.g. 0.305 for *MYC*, 0.685 for *CDK4*, 0.778 for *MDM2*, 0.701 for *KRAS*, 0.414 for *MTOR*) and higher than 1 when TSG are knocked out (1.411 for *TP53*, 1.792 for *PTEN*, 1.170 for *RB1*, 1.227 for *CDKN1A*, 1.136 for *BAX*), all values being highly significantly different from those of *AGR2* and *AGR3* (*p* values ranging from 10^−9^ to 0). As a consequence, *AGR2* and *AGR3* appear in this respect as “neutral” genes, whose knock-outs have very moderate influence on cell fitness. However, when building a heatmap with normalised ranked values of cell fitness alterations induced by ten major oncogenes and ten major TSG (Figure 8), a good segregation between oncogenes and TSG clearly appears, with *AGR2* and *AGR3* segregating together among oncogenes. We evaluated also the correlations that could exist between the extinction patterns of *AGR2* and *AGR3* to those of other genes (Supplemental Table 5). It appeared that, among the 60 genes presenting a pattern of extinction significantly correlated (down to 10^−6^) with that of *AGR2*, 44 are localised on chromosome arm 7p, indicating a topologic rather than a functional relationship. Whereas there was no oncogene or TSG among the genes located on chromosome arm 7p, there were three oncogenes (*KLF5, TCF7L2* and *CTNNB1*) and one TSG (*SOX9*) located in other chromosome arms, all presenting an extinction pattern similar to that of *AGR2* among the CCLE collection (positive correlation) and playing a role in transcription. *AGR3* displayed a distinct pattern of gene extinction, with only 9 genes not located on 7p chromosome arm out of 105 whose extinction pattern was correlated with that of this gene, which does not bring information on functional relationship.

**Figure 8.**
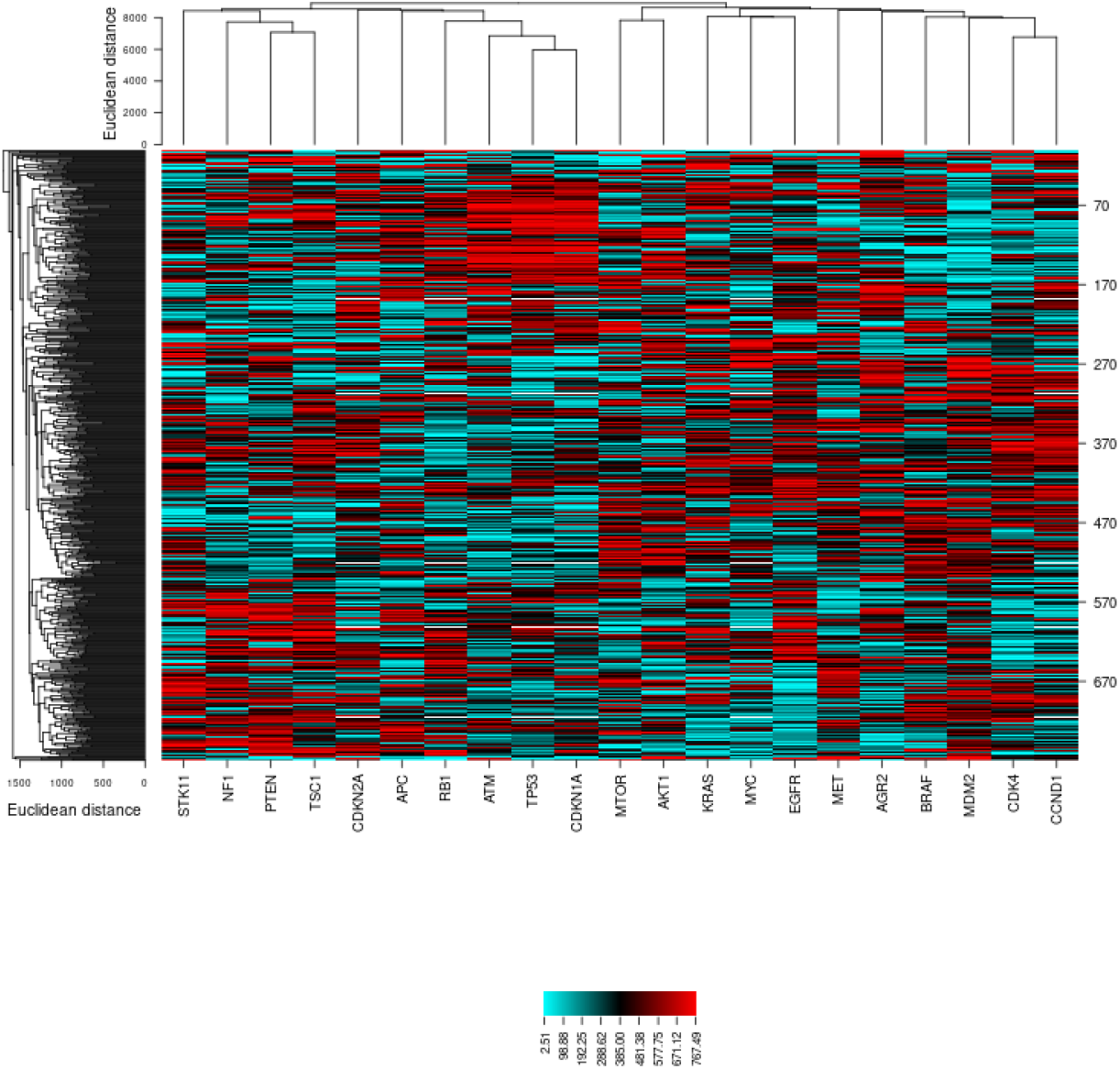
Clustering of cell fitness alterations in various oncogenes and TSG. Clustering was performed using CIMminer on the NCI Genomics and Pharmacology Facility (https://discover.nci.nih.gov/cimminer/oneMatrix.do). Fitness values were downloaded and normalised by ranking before building the heatmap.

## Discussion

The question underlying the development of this work is whether *AGR2* and *AGR3* can be considered as playing a major role in oncogenesis and progression of cancers; in other terms, whether they can be considered as oncogenes and/or TSG. Known polymorphisms in *AGR2* and *AGR3* sequences as well as variations encountered at a known polymorphic site are not likely to confer oncogenic properties to AGR2 or AGR3 proteins. Only five SNV in *AGR2* and six in *AGR3* sequence deserve some attention: those which are supposed to result in a truncated or different protein (nonsense, frameshift variations). These variations are not recurrent and cannot be considered as oncogenic variations since the tumours and cell lines bearing these variations do not behave differently than the other ones in terms of *AGR2*/*3* gene expression.

Similarly, the *AGR2*/*3* copy number variations encountered in TCGA did not seem to affect *AGR2*/*3* gene expression. However, we observed a significant negative correlation between *AGR2* expression and chromosome 7p deletions in BRCA, which could be expected since this is the chromosome location of *AGR2*/*3*. In the CCLE, when the whole set of cell lines was taken in consideration, there was a significant correlation for both genes between *AGR2* gene copy number and expression. When ranking the copy number values from highest to lowest values, there was no preferential contribution of the cancer types to presenting high or low *AGR2*/*3* copy numbers. The overall conclusion of these explorations of *AGR2* and *AGR3* genomic variations in tumours and cancer cell lines is that it is quite unlikely that they could behave as *bona fide* oncogenes or TSG.

The associations we noticed between *AGR2* gene expression and that of a large series of genes reveal in contrast several important features in relation to oncogenesis and cancer progression. A common general feature is the fact that both genes appear as epithelial markers, in TCGA different cancer types as well as in the whole set of CCLE cell lines and in cell lines of different cancer types. In addition, there was a negative correlation between *AGR2* expression and that of the main transcription factors of epithelial-to-mesenchymal transition. Another point of interest is that some of the known partners of AGR2 and AGR3 proteins are co-expressed with them, but this is not a general feature, and concerns the different cancer types in a specific way, with the exception of mucins whose expression appears to be strongly positively correlated to that of *AGR2*/*3* in all cancer types, in agreement with their known functional association.

It appears from our explorations that AGR2 and AGR3 are connected to the cancer phenotype. In clinical samples as well as in CCLE cancer cell lines, the presence of oncogenic mutations and CNVs in various driver genes is associated with variations in *AGR2*/*3* expression, depending both of the cancer gene and the tumour type.

*AGR2* gene extinction in CRISPR screens of the CCLE is followed by a mitigate, low amplitude consequence on cell survival and proliferation, with a null average value, whereas oncogene or TSG extinction is followed by significant effects, either in favour (oncogenes) or to the detriment (TSG) of cell fitness. *AGR2* gene extinction pattern appears to be correlated to that of several cancer genes, reinforcing the attribution of a participation of this protein in cancer phenotypes.

It is commonly assumed that somatic mutations drive the multi-step tumour development process. Although *AGR2* and *AGR3* genes present no recurrent mutations, both proteins are often overexpressed, have non-canonical localisations (extracellular, cytosol) and are associated to different tumour processes such as differentiation, proliferation, migration, invasion, metastasis, in almost all epithelial cancer types. Cancer follows an evolutionary trajectory, characterised by stepwise acquisition of mutations that allow the tumour cells to increase their fitness, from the precancer lesion to tumour metastasis. However, the non-genetic gain-of-functions alterations, acquired by overexpression and non-canonical localisations of AGR2 and AGR3 proteins, may be pivotal for tumour development and progression.

Thus, AGR2 and AGR3 proteins appear as common non-genetic evolutionary factors in the process of human tumorigenesis. Complex and dynamic adaptation mechanisms and evolutionary processes take place during the process of human epithelial tumorigenesis (tumour initiation, development and progression). Although cancer has been considered mainly, for decades, as a process governed by genetic mechanisms, it is becoming clearer that non-genetic mechanisms may also play an important role in cancer progression. Tumours are constantly evolving, displaying highly variable patterns resulting in extremely complex genetic and non-genetic phenotypic diversification. Therefore, when dealing with such a complex system that is barely understood, common hallmarks are rare. Thus, it is of crucial importance to identify and investigate the functional role of novel unexpected common hallmarks that will undoubtedly aid the development of therapeutic approaches. Overexpression and non-canonical localisations of AGR2 and AGR3 may reflect a non-genetic evolutive process, which is indeed a common feature in human epithelial tumorigenesis. We believe that further in-depth functional studies of cancer development from an AGR2/3 expression and localisation perspective may enable us to progress in the understanding of the epithelial cancer evolutionary framework, which might result in the discovery of new original therapeutic perspectives.

## Acknowledgements

We gratefully acknowledge the members from ARTiSt group for their critical remarks. FD was supported by grant from the “*Fondation ARC pour la recherche sur le cancer*”, FD and IV from the “*Site de recherche intégrée sur le cancer de Bordeaux*” (SIRIC Brio). This work has been supported by grants from the “*Région Nouvelle-Aquitaine*” (DF and FD), by the “*Agence Nationale de la Recherche”* (ANR) (DF), and by the “Ligue contre le cancer Gironde” (FD). This work was also funded by grants from the “*Institut National du Cancer*” (INCa, PLBIO), “*Fondation pour la Recherche Médicale*” (FRM, DEQ20180339169), and “*Agence Nationale de la Recherche*” (ANR; ERAAT) to EC.

---

**Supplemental Table 1.**
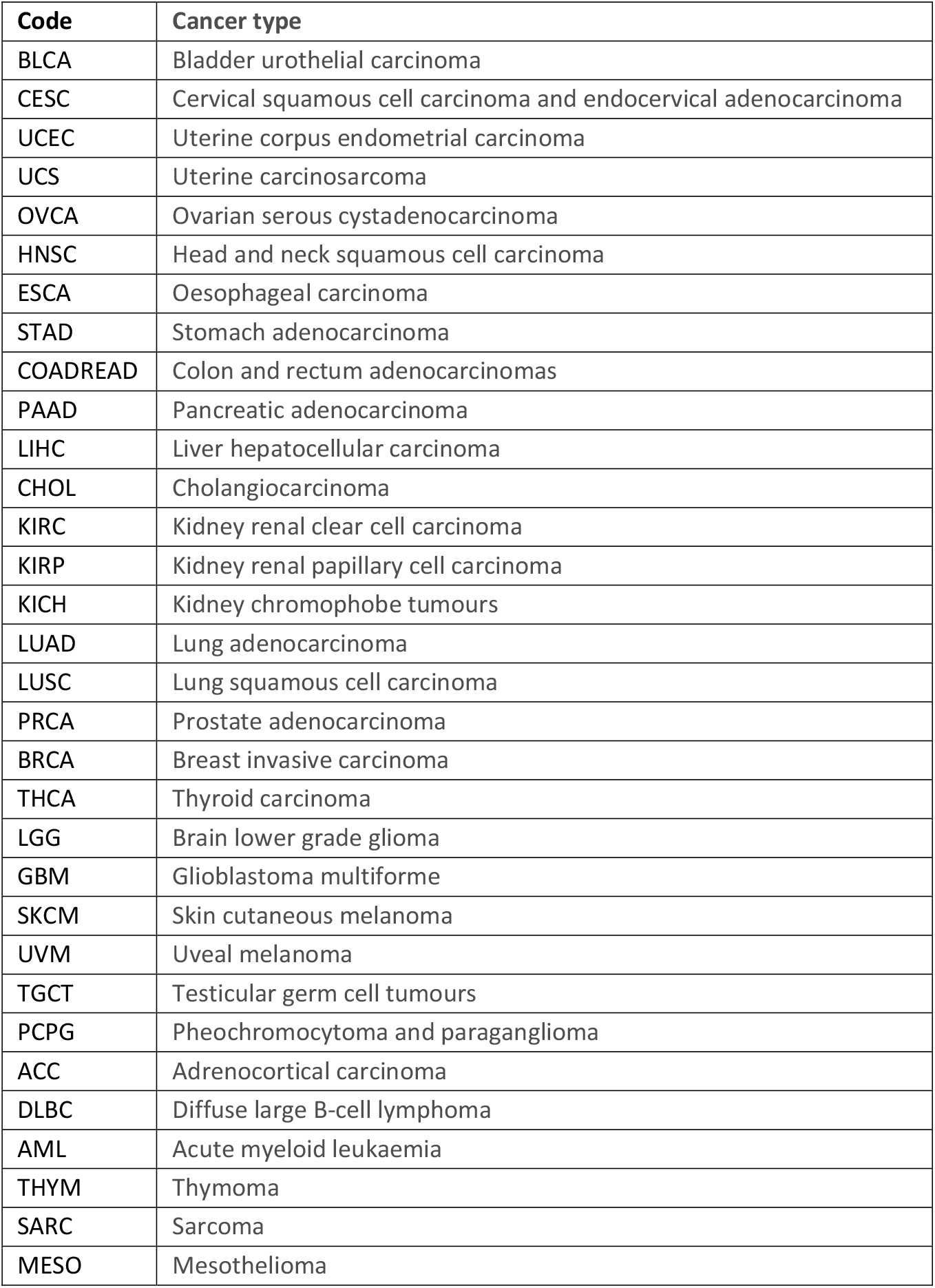
Nomenclature of the cancer types included in TCGA.

**Supplemental Table 2A.** Prevalence of AGR2 genome alterations in 9,888 tumour samples of TCGA. Out of the 10,947 samples in TCGA PanCancer Atlas, 9,888 were suited for genomic analyses (mutation and CNA data).

|  | Nb total | Mutations |  | Amplifications |  | Homoz. deletions |  |
| --- | --- | --- | --- | --- | --- | --- | --- |
|  |  | Nb | % | Nb | % | Nb | % |
| BLCA | 402 | 2 | 0.49 | 21 | 5.11 | 1 | 0.24 |
| CESC | 278 | 2 | 0.67 | 2 | 0.67 |  |  |
| UCEC | 509 | 11 | 2.08 | 7 | 1.32 | 2 | 0.38 |
| UCS | 56 |  |  | 2 | 3.51 |  |  |
| OVCA | 398 |  |  | 8 | 1.37 | 6 | 1.03 |
| HNSC | 496 |  |  | 10 | 1.91 |  |  |
| ESCA | 182 |  |  | 10 | 5.49 |  |  |
| STAD | 434 |  |  | 13 | 2.95 |  |  |
| COADREAD | 524 | 2 | 0.34 | 2 | 0.34 |  |  |
| PAAD | 175 | 1 | 0.54 | 1 | 0.54 |  |  |
| LIHC | 353 | 1 | 0.27 |  |  |  |  |
| CHOL | 36 |  |  |  |  |  |  |
| KIRC | 354 |  |  | 3 | 0.59 |  |  |
| KIRP | 274 | 1 | 0.35 |  |  |  |  |
| KICH | 65 |  |  |  |  |  |  |
| LUAD | 507 | 1 | 0.18 | 18 | 3.18 | 2 | 0.35 |
| LUSC | 469 |  |  | 7 | 1.44 |  |  |
| PRCA | 489 | 1 | 0.20 | 6 | 1.21 | 2 | 0.40 |
| BRCA | 994 |  |  | 7 | 0.65 |  |  |
| THCA | 482 | 1 | 0.20 | 1 | 0.20 |  |  |
| LGG | 511 |  |  | 3 | 0.58 |  |  |
| GBM | 378 | 1 | 0.17 | 5 | 0.84 |  |  |
| SKCM | 363 | 7 | 1.56 | 7 | 1.56 |  |  |
| UVM | 80 |  |  |  |  |  |  |
| TGCT | 144 |  |  | 2 | 1.34 |  |  |
| PCPG | 161 |  |  | 1 | 0.56 |  |  |
| ACC | 89 |  |  | 3 | 3.26 |  |  |
| DLBC | 37 |  |  |  |  | 1 | 2.08 |
| AML | 190 |  |  |  |  |  |  |
| THYM | 123 |  |  |  |  |  |  |
| SARC | 253 | 1 | 0.39 | 7 | 2.75 |  |  |
| MESO | 82 |  |  |  |  |  |  |
| **Total** | **9,888** | **32** | **0.3** | **146** | **1.41** | **14** | **0.13** |

**Supplemental Table 2B.** Prevalence of AGR3 genome alterations in 9,888 tumour samples of TCGA. Out of the 10,947 samples in TCGA PanCancer Atlas, 9,888 were suited for genomic analyses (mutation and CNA data).

|  | Nb total | Mutations |  | Amplifications |  | Homoz. deletions |  |
| --- | --- | --- | --- | --- | --- | --- | --- |
|  |  | Nb | % | Nb | % | Nb | % |
| BLCA | 402 | 6 | 1.49 | 21 | 5.22 | 1 | 0.25 |
| CESC | 278 | 2 | 0.72 | 2 | 0.72 |  |  |
| UCEC | 509 | 6 | 1.18 | 9 | 1.77 | 2 | 0.39 |
| UCS | 56 |  |  | 2 | 3.57 |  |  |
| OVCA | 398 |  |  | 8 | 2.01 | 6 | 1.51 |
| HNSC | 496 | 1 | 0.20 | 10 | 2.02 |  |  |
| ESCA | 182 | 1 | 0.55 | 8 | 4.40 |  |  |
| STAD | 434 | 5 | 1.15 | 13 | 3.00 | 1 | 0.23 |
| COADREAD | 524 | 1 | 0.19 | 2 | 0.38 |  |  |
| PAAD | 175 |  |  | 1 | 0.57 |  |  |
| LIHC | 353 | 1 | 0.28 |  |  |  |  |
| CHOL | 36 |  |  |  |  |  |  |
| KIRC | 354 |  |  | 4 | 1.13 |  |  |
| KIRP | 274 |  |  |  |  |  |  |
| KICH | 65 |  |  |  |  |  |  |
| LUAD | 507 | 2 | 0.39 | 18 | 3.55 | 2 | 0.39 |
| LUSC | 469 | 1 | 0.21 | 8 | 1.71 |  |  |
| PRCA | 489 | 1 | 0.20 | 6 | 1.23 | 2 | 0.41 |
| BRCA | 994 |  |  | 7 | 0.70 |  |  |
| THCA | 482 |  |  | 1 | 0.21 |  |  |
| LGG | 511 | 1 | 0.20 | 2 | 0.39 |  |  |
| GBM | 378 |  |  | 4 | 1.06 |  |  |
| SKCM | 363 | 6 | 1.65 | 7 | 1.93 |  |  |
| UVM | 80 |  |  |  |  |  |  |
| TGCT | 144 |  |  | 2 | 1.39 |  |  |
| PCPG | 161 |  |  | 1 | 0.62 |  |  |
| ACC | 89 | 1 | 1.12 | 3 | 3.37 |  |  |
| DLBC | 37 |  |  |  |  | 1 | 2.70 |
| AML | 190 |  |  |  |  |  |  |
| THYM | 123 |  |  |  |  |  |  |
| SARC | 253 |  |  | 6 | 2.77 |  |  |
| MESO | 82 |  |  |  |  |  |  |
| **Total** | **9,888** | **35** | **0.35** | **145** | **1.47** | **15** | **0.15** |

**Supplemental Table 3.** List of oncogenes and TSG whose mutations are positively or negatively correlated with *AGR2* and *AGR3* expressions (*p* values ranking, down to 10^−4^) in the whole set of cell lines of the CCLE collection. Oncogenes are spotted in red and TSG in blue. Oncogenes and TSG were extracted among a list of 568 cancer driver genes from the Integrative OncoGenomics platform (https://www.intogen.org/search). It should be underlined that some genes, especially some involved in transcriptional control, may behave as oncogenes or TSG according to the context.

| *AGR2* |  |  |  | *AGR3* |  |  |  |
| --- | --- | --- | --- | --- | --- | --- | --- |
| Gene | Cytoband | *r* | *p* | Gene | Cytoband | *r* | *p* |
| *KRAS* | 12p12.1 | 0.237 | 6.79×10<sup>-13</sup> | *APC* | 5q21-q22 | 0.246 | 8.22×10<sup>-14</sup> |
| *SMAD4* | 18q21.1 | 0.220 | 2.68×10<sup>-11</sup> | *ACVR2A* | 2q22.3 | 0.231 | 2.58×10<sup>-12</sup> |
| *PIK3CA* | 3q26.3 | 0.197 | 2.76×10<sup>-9</sup> | *SMAD4* | 18q21.1 | 0.222 | 1.70×10<sup>-11</sup> |
| *TGFBR1* | 9q22 | 0.188 | 1.31×10<sup>-8</sup> | *SOST* | 17q11.2 | 0.209 | 2.68×10<sup>-10</sup> |
| *APC* | 5q21-q22 | 0.187 | 1.55×10<sup>-8</sup> | *MAP3K4* | 6q26 | 0.197 | 2.76×10<sup>-9</sup> |
| *RHOA* | 3p21.3 | 0.178 | 7.89×10<sup>-8</sup> | *NIPBL* | 5p13.2 | 0.190 | 9.25×10<sup>-9</sup> |
| *ACVR2A* | 2q22.3 | 0.177 | 9.72×10<sup>-8</sup> | *CDH1* | 16q22.1 | 0.175 | 1.24×10<sup>-7</sup> |
| *CDH1* | 16q22.1 | 0.173 | 1.80×10<sup>-7</sup> | *FBXW7* | 4q31.3 | 0.175 | 1.38×10<sup>-7</sup> |
| *PARP1* | 1q41-q42 | 0.149 | 7.02×10<sup>-6</sup> | *PARP1* | 1q41-q42 | 0.172 | 2.27×10<sup>-7</sup> |
| *CDC25C* | 5q31 | 0.148 | 8.08×10<sup>-6</sup> | *NTRK3* | 15q25 | 0.167 | 4.64×10<sup>-7</sup> |
| *MAP3K4* | 6q26 | 0.139 | 2.80×10<sup>-5</sup> | *ATR* | 3q23 | 0.165 | 6.59×10<sup>-7</sup> |
| *ACVR2B* | 3p22 | 0.139 | 3.12×10<sup>-5</sup> | *RHOA* | 3p21.3 | 0.162 | 1.12×10<sup>-6</sup> |
| *FBXW7* | 4q31.3 | 0.137 | 3.78×10<sup>-5</sup> | *MGA* | 15q14 | 0.159 | 1.69×10<sup>-6</sup> |
| *SOST* | 17q11.2 | 0.136 | 4.34×10<sup>-5</sup> | *PIK3CA* | 3q26.3 | 0.157 | 2.33×10<sup>-6</sup> |
| *TGFBR2* | 3p22 | 0.135 | 5.06×10<sup>-5</sup> | *CDC25C* | 5q31 | 0.155 | 2.91×10<sup>-6</sup> |
| *NIPBL* | 5p13.2 | 0.130 | 9.42×10<sup>-5</sup> | *TGFBR2* | 3p22 | 0.155 | 3.12×10<sup>-6</sup> |
|  |  |  |  | *HDAC1* | 1p34 | 0.154 | 3.61×10<sup>-6</sup> |
|  |  |  |  | *MDM2* | 12q14.3-q15 | 0.154 | 3.81×10<sup>-6</sup> |
|  |  |  |  | *SMAD2* | 18q21.1 | 0.147 | 1.02×10<sup>-5</sup> |
|  |  |  |  | *CDC73* | 1q25 | 0.146 | 1.20×10<sup>-5</sup> |
|  |  |  |  | *AXIN1* | 16p13.3 | 0.145 | 1.25×10<sup>-5</sup> |
|  |  |  |  | *RAD50* | 5q31 | 0.145 | 1.26×10<sup>-5</sup> |
|  |  |  |  | *BCL6* | 3q27 | 0.145 | 1.27×10<sup>-5</sup> |
|  |  |  |  | *KDM5A* | 12p11 | 0.145 | 1.33×10<sup>-5</sup> |
|  |  |  |  | *CARD11* | 7p22 | 0.141 | 2.13×10<sup>-5</sup> |
|  |  |  |  | *SPEN* | 1p36 | 0.139 | 3.10×10<sup>-5</sup> |
|  |  |  |  | *CNTN6* | 3p26-p25 | 0.137 | 3.74×10<sup>-5</sup> |
|  |  |  |  | *RAF1* | 3p25 | 0.136 | 4.47×10<sup>-5</sup> |
|  |  |  |  | *AKT1* | 14q32.32 | 0.135 | 5.15×10<sup>-5</sup> |
|  |  |  |  | *AKT3* | 1q44 | 0.135 | 5.16×10<sup>-5</sup> |
|  |  |  |  | *MAP2K4* | 17p12 | 0.134 | 5.50×10<sup>-5</sup> |
|  |  |  |  | *BAX* | 19q13.3-q13.4 | 0.133 | 6.30×10<sup>-5</sup> |
|  |  |  |  | *MAPK1* | 22q11.21 | 0.133 | 6.50×10<sup>-5</sup> |
|  |  |  |  | *ATM* | 11q22-q23 | 0.131 | 8.06×10<sup>-5</sup> |
|  |  |  |  | *RPL22* | 1p36.31 | 0.130 | 9.60×10<sup>-5</sup> |

**Supplemental Table 4A.**
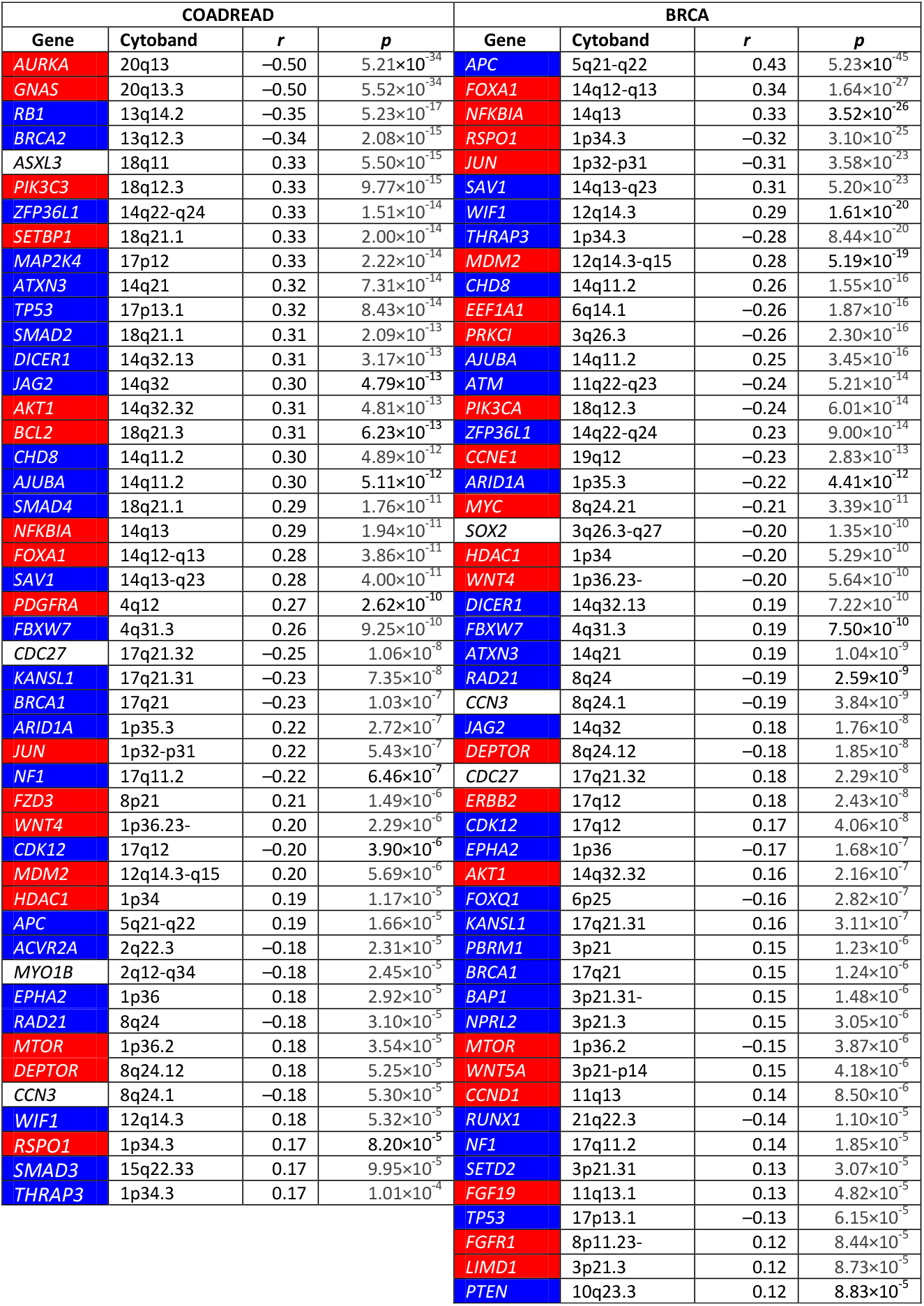
List of oncogenes and TSG whose copy numbers are positively or negatively correlated with *AGR2* expression in the COADREAD and BRCA samples of TCGA (*p* value ranking, down to p < 10^−4^) Oncogenes are spotted in red and TSG in blue. Oncogenes and TSG were extracted among a list of 568 cancer driver genes from the Integrative OncoGenomics platform (https://www.intogen.org/search). It should be underlined that some genes, especially some involved in transcriptional control, may behave as oncogenes or TSG according to the context.

**Supplemental Table 4B.**
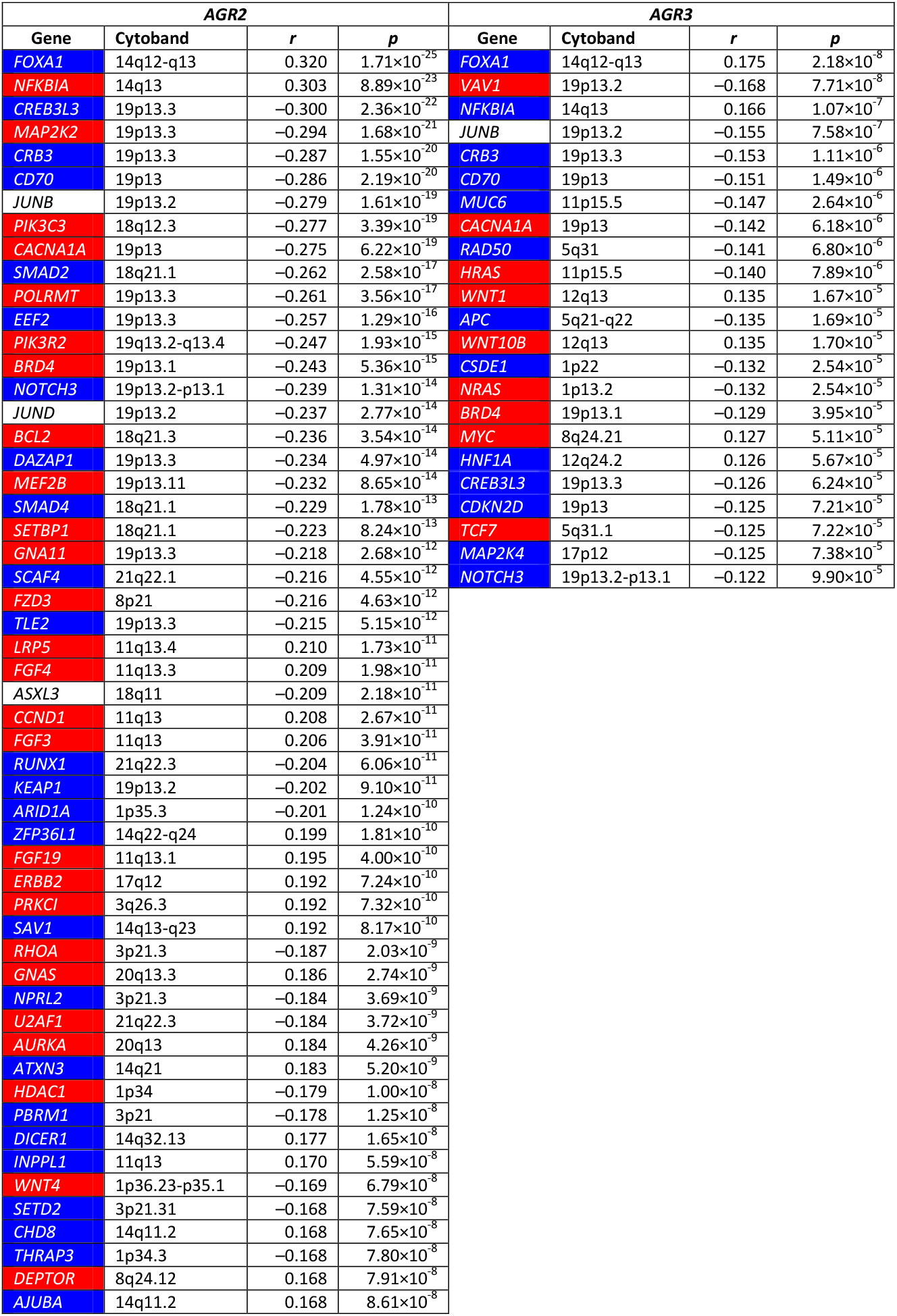
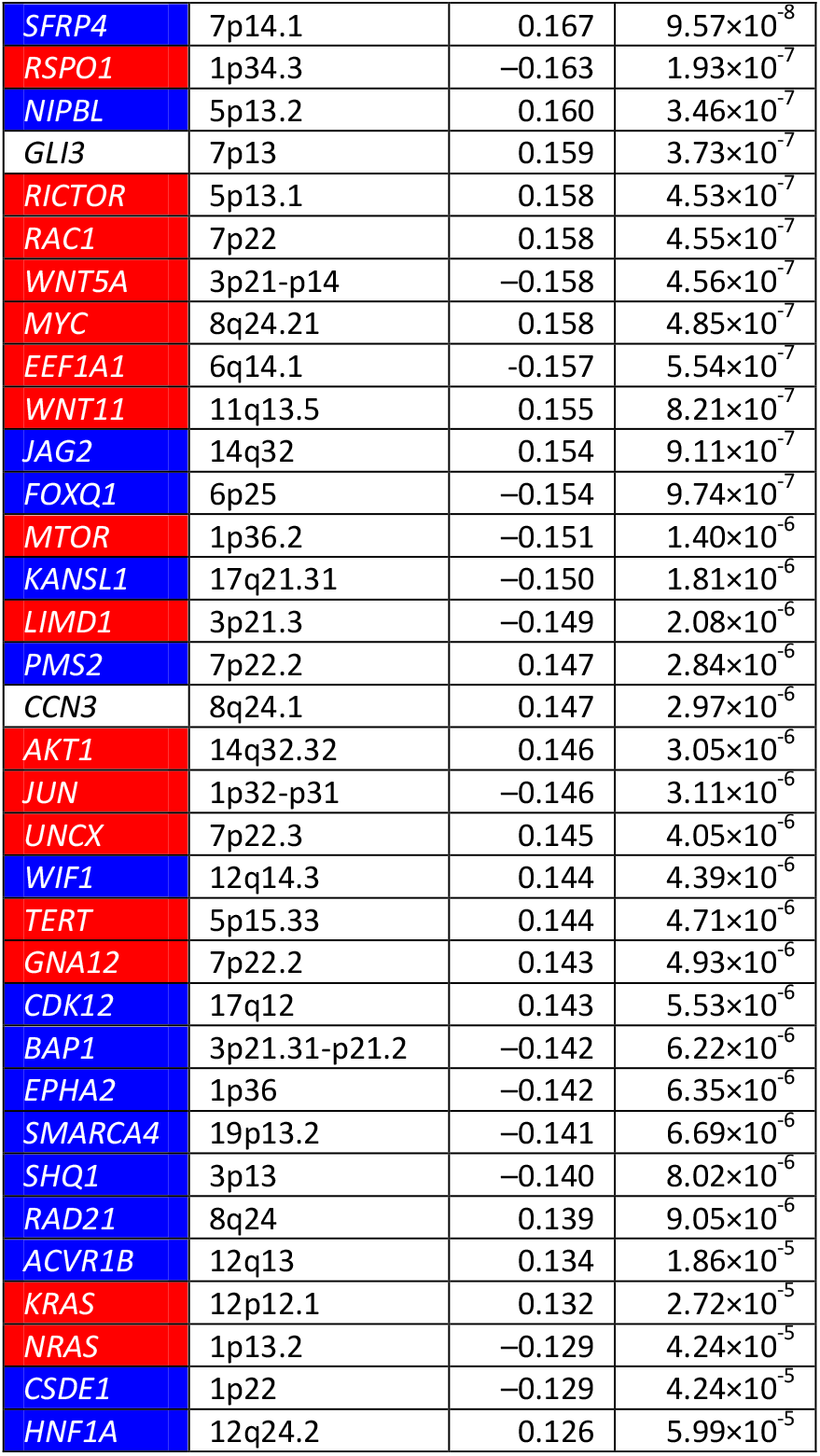
List of oncogenes and TSG whose copy numbers are positively or negatively correlated with *AGR2* and *AGR3* expressions in the whole set of cell lines of the CCLE collection (*p* value ranking, down to p < 10^−4^) Oncogenes are spotted in red and TSG in blue. Oncogenes and TSG were extracted among a list of 568 cancer driver genes from the Integrative OncoGenomics platform (https://www.intogen.org/search). It should be underlined that some genes, especially some involved in transcriptional control, may behave as oncogenes or TSG according to the context.

| *AGR2* |  |  |  | *AGR3* |  |  |  |
| --- | --- | --- | --- | --- | --- | --- | --- |
| Gene | Cytoband | *r* | *p* | Gene | Cytoband | *r* | *p* |
| *FOXA1* | 14q12-q13 | 0.320 | $1.71 \times 10^{-25}$ | *FOXA1* | 14q12-q13 | 0.175 | $2.18 \times 10^{-8}$ |
| *NFKBIA* | 14q13 | 0.303 | $8.89 \times 10^{-23}$ | *VAV1* | 19p13.2 | -0.168 | $7.71 \times 10^{-8}$ |
| *CREB3L3* | 19p13.3 | -0.300 | $2.36 \times 10^{-22}$ | *NFKBIA* | 14q13 | 0.166 | $1.07 \times 10^{-7}$ |
| *MAP2K2* | 19p13.3 | -0.294 | $1.68 \times 10^{-21}$ | *JUNB* | 19p13.2 | -0.155 | $7.58 \times 10^{-7}$ |
| *CRB3* | 19p13.3 | -0.287 | $1.55 \times 10^{-20}$ | *CRB3* | 19p13.3 | -0.153 | $1.11 \times 10^{-6}$ |
| *CD70* | 19p13 | -0.286 | $2.19 \times 10^{-20}$ | *CD70* | 19p13 | -0.151 | $1.49 \times 10^{-6}$ |
| *JUNB* | 19p13.2 | -0.279 | $1.61 \times 10^{-19}$ | *MUC6* | 11p15.5 | -0.147 | $2.64 \times 10^{-6}$ |
| *PIK3C3* | 18q12.3 | -0.277 | $3.39 \times 10^{-19}$ | *CACNA1A* | 19p13 | -0.142 | $6.18 \times 10^{-6}$ |
| *CACNA1A* | 19p13 | -0.275 | $6.22 \times 10^{-19}$ | *RAD50* | 5q31 | -0.141 | $6.80 \times 10^{-6}$ |
| *SMAD2* | 18q21.1 | -0.262 | $2.58 \times 10^{-17}$ | *HRAS* | 11p15.5 | -0.140 | $7.89 \times 10^{-6}$ |
| *POLRMT* | 19p13.3 | -0.261 | $3.56 \times 10^{-17}$ | *WNT1* | 12q13 | 0.135 | $1.67 \times 10^{-5}$ |
| *EEF2* | 19p13.3 | -0.257 | $1.29 \times 10^{-16}$ | *APC* | 5q21-q22 | -0.135 | $1.69 \times 10^{-5}$ |
| *PIK3R2* | 19q13.2-q13.4 | -0.247 | $1.93 \times 10^{-15}$ | *WNT10B* | 12q13 | 0.135 | $1.70 \times 10^{-5}$ |
| *BRD4* | 19p13.1 | -0.243 | $5.36 \times 10^{-15}$ | *CSDE1* | 1p22 | -0.132 | $2.54 \times 10^{-5}$ |
| *NOTCH3* | 19p13.2-p13.1 | -0.239 | $1.31 \times 10^{-14}$ | *NRAS* | 1p13.2 | -0.132 | $2.54 \times 10^{-5}$ |
| *JUND* | 19p13.2 | -0.237 | $2.77 \times 10^{-14}$ | *BRD4* | 19p13.1 | -0.129 | $3.95 \times 10^{-5}$ |
| *BCL2* | 18q21.3 | -0.236 | $3.54 \times 10^{-14}$ | *MYC* | 8q24.21 | 0.127 | $5.11 \times 10^{-5}$ |
| *DAZAP1* | 19p13.3 | -0.234 | $4.97 \times 10^{-14}$ | *HNF1A* | 12q24.2 | 0.126 | $5.67 \times 10^{-5}$ |
| *MEF2B* | 19p13.11 | -0.232 | $8.65 \times 10^{-14}$ | *CREB3L3* | 19p13.3 | -0.126 | $6.24 \times 10^{-5}$ |
| *SMAD4* | 18q21.1 | -0.229 | $1.78 \times 10^{-13}$ | *CDKN2D* | 19p13 | -0.125 | $7.21 \times 10^{-5}$ |
| *SETBP1* | 18q21.1 | -0.223 | $8.24 \times 10^{-13}$ | *TCF7* | 5q31.1 | -0.125 | $7.22 \times 10^{-5}$ |
| *GNA11* | 19p13.3 | -0.218 | $2.68 \times 10^{-12}$ | *MAP2K4* | 17p12 | -0.125 | $7.38 \times 10^{-5}$ |
| *SCAF4* | 21q22.1 | -0.216 | $4.55 \times 10^{-12}$ | *NOTCH3* | 19p13.2-p13.1 | -0.122 | $9.90 \times 10^{-5}$ |
| *FZD3* | 8p21 | -0.216 | $4.63 \times 10^{-12}$ | | | | |
| *TLE2* | 19p13.3 | -0.215 | $5.15 \times 10^{-12}$ | | | | |
| *LRP5* | 11q13.4 | 0.210 | $1.73 \times 10^{-11}$ | | | | |
| *FGF4* | 11q13.3 | 0.209 | $1.98 \times 10^{-11}$ | | | | |
| *ASXL3* | 18q11 | -0.209 | $2.18 \times 10^{-11}$ | | | | |
| *CCND1* | 11q13 | 0.208 | $2.67 \times 10^{-11}$ | | | | |
| *FGF3* | 11q13 | 0.206 | $3.91 \times 10^{-11}$ | | | | |
| *RUNX1* | 21q22.3 | -0.204 | $6.06 \times 10^{-11}$ | | | | |
| *KEAP1* | 19p13.2 | -0.202 | $9.10 \times 10^{-11}$ | | | | |
| *ARID1A* | 1p35.3 | -0.201 | $1.24 \times 10^{-10}$ | | | | |
| *ZFP36L1* | 14q22-q24 | 0.199 | $1.81 \times 10^{-10}$ | | | | |
| *FGF19* | 11q13.1 | 0.195 | $4.00 \times 10^{-10}$ | | | | |
| *ERBB2* | 17q12 | 0.192 | $7.24 \times 10^{-10}$ | | | | |
| *PRKCI* | 3q26.3 | 0.192 | $7.32 \times 10^{-10}$ | | | | |
| *SAV1* | 14q13-q23 | 0.192 | $8.17 \times 10^{-10}$ | | | | |
| *RHOA* | 3p21.3 | -0.187 | $2.03 \times 10^{-9}$ | | | | |
| *GNAS* | 20q13.3 | 0.186 | $2.74 \times 10^{-9}$ | | | | |
| *NPRL2* | 3p21.3 | -0.184 | $3.69 \times 10^{-9}$ | | | | |
| *U2AF1* | 21q22.3 | -0.184 | $3.72 \times 10^{-9}$ | | | | |
| *AURKA* | 20q13 | 0.184 | $4.26 \times 10^{-9}$ | | | | |
| *ATXN3* | 14q21 | 0.183 | $5.20 \times 10^{-9}$ | | | | |
| *HDAC1* | 1p34 | -0.179 | $1.00 \times 10^{-8}$ | | | | |
| *PBRM1* | 3p21 | -0.178 | $1.25 \times 10^{-8}$ | | | | |
| *DICER1* | 14q32.13 | 0.177 | $1.65 \times 10^{-8}$ | | | | |
| *INPPL1* | 11q13 | 0.170 | $5.59 \times 10^{-8}$ | | | | |
| *WNT4* | 1p36.23-p35.1 | -0.169 | $6.79 \times 10^{-8}$ | | | | |
| *SETD2* | 3p21.31 | -0.168 | $7.59 \times 10^{-8}$ | | | | |
| *CHD8* | 14q11.2 | 0.168 | $7.65 \times 10^{-8}$ | | | | |
| *THRAP3* | 1p34.3 | -0.168 | $7.80 \times 10^{-8}$ | | | | |
| *DEPTOR* | 8q24.12 | 0.168 | $7.91 \times 10^{-8}$ | | | | |
| *AJUBA* | 14q11.2 | 0.168 | $8.61 \times 10^{-8}$ | | | | |

|  |  |  |  |
| --- | --- | --- | --- |
| **SFRP4** | 7p14.1 | 0.167 | $9.57 \times 10^{-8}$ |
| **RSPO1** | 1p34.3 | -0.163 | $1.93 \times 10^{-7}$ |
| **NIPBL** | 5p13.2 | 0.160 | $3.46 \times 10^{-7}$ |
| **GLI3** | 7p13 | 0.159 | $3.73 \times 10^{-7}$ |
| **RICTOR** | 5p13.1 | 0.158 | $4.53 \times 10^{-7}$ |
| **RAC1** | 7p22 | 0.158 | $4.55 \times 10^{-7}$ |
| **WNT5A** | 3p21-p14 | -0.158 | $4.56 \times 10^{-7}$ |
| **MYC** | 8q24.21 | 0.158 | $4.85 \times 10^{-7}$ |
| **EEF1A1** | 6q14.1 | -0.157 | $5.54 \times 10^{-7}$ |
| **WNT11** | 11q13.5 | 0.155 | $8.21 \times 10^{-7}$ |
| **JAG2** | 14q32 | 0.154 | $9.11 \times 10^{-7}$ |
| **FOXQ1** | 6p25 | -0.154 | $9.74 \times 10^{-7}$ |
| **MTOR** | 1p36.2 | -0.151 | $1.40 \times 10^{-6}$ |
| **KANSL1** | 17q21.31 | -0.150 | $1.81 \times 10^{-6}$ |
| **LIMD1** | 3p21.3 | -0.149 | $2.08 \times 10^{-6}$ |
| **PMS2** | 7p22.2 | 0.147 | $2.84 \times 10^{-6}$ |
| **CCN3** | 8q24.1 | 0.147 | $2.97 \times 10^{-6}$ |
| **AKT1** | 14q32.32 | 0.146 | $3.05 \times 10^{-6}$ |
| **JUN** | 1p32-p31 | -0.146 | $3.11 \times 10^{-6}$ |
| **UNCX** | 7p22.3 | 0.145 | $4.05 \times 10^{-6}$ |
| **WIF1** | 12q14.3 | 0.144 | $4.39 \times 10^{-6}$ |
| **TERT** | 5p15.33 | 0.144 | $4.71 \times 10^{-6}$ |
| **GNA12** | 7p22.2 | 0.143 | $4.93 \times 10^{-6}$ |
| **CDK12** | 17q12 | 0.143 | $5.53 \times 10^{-6}$ |
| **BAP1** | 3p21.31-p21.2 | -0.142 | $6.22 \times 10^{-6}$ |
| **EPHA2** | 1p36 | -0.142 | $6.35 \times 10^{-6}$ |
| **SMARCA4** | 19p13.2 | -0.141 | $6.69 \times 10^{-6}$ |
| **SHQ1** | 3p13 | -0.140 | $8.02 \times 10^{-6}$ |
| **RAD21** | 8q24 | 0.139 | $9.05 \times 10^{-6}$ |
| **ACVR1B** | 12q13 | 0.134 | $1.86 \times 10^{-5}$ |
| **KRAS** | 12p12.1 | 0.132 | $2.72 \times 10^{-5}$ |
| **NRAS** | 1p13.2 | -0.129 | $4.24 \times 10^{-5}$ |
| **CSDE1** | 1p22 | -0.129 | $4.24 \times 10^{-5}$ |
| **HNF1A** | 12q24.2 | 0.126 | $5.99 \times 10^{-5}$ |

**Supplemental Table 5.**
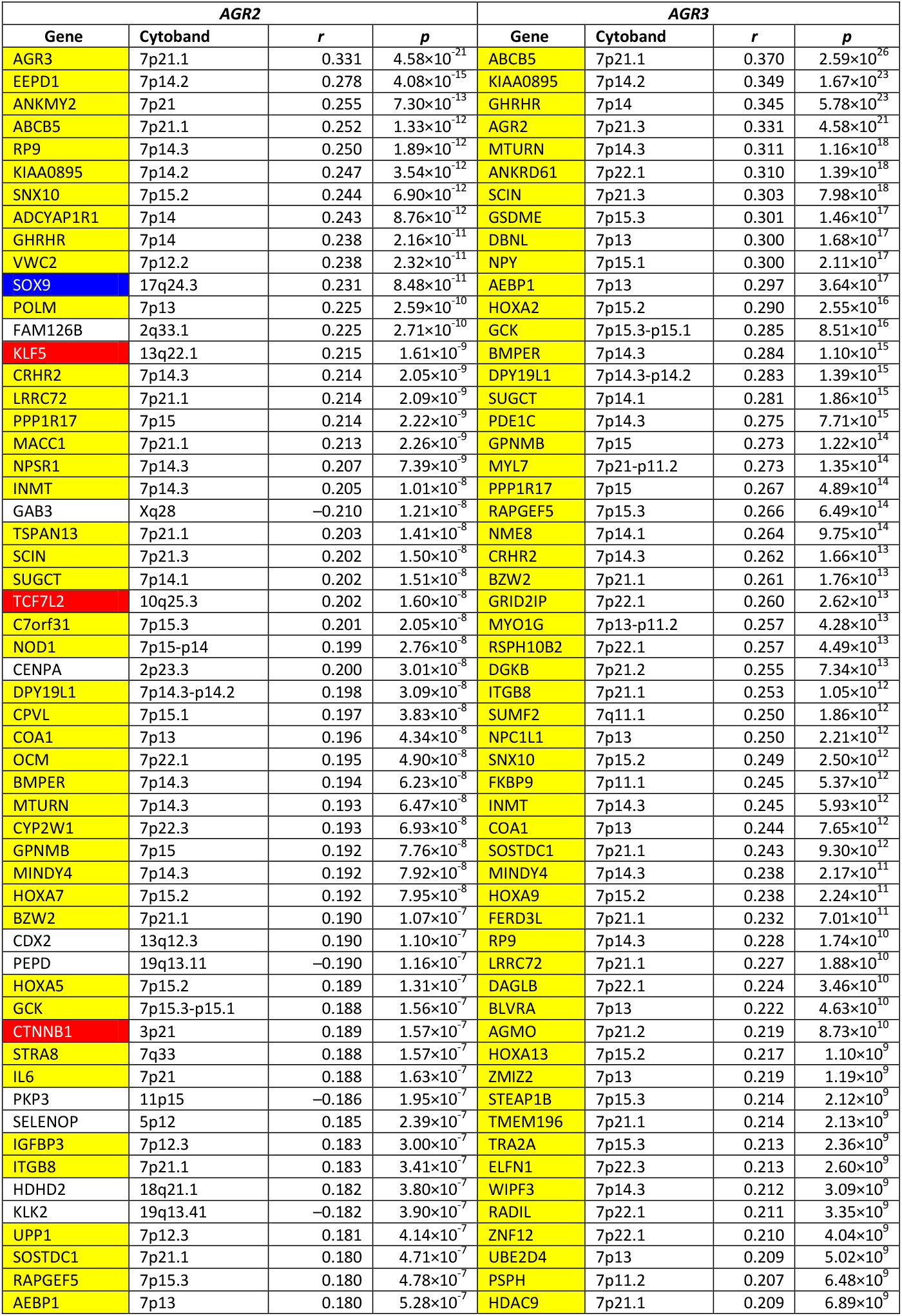
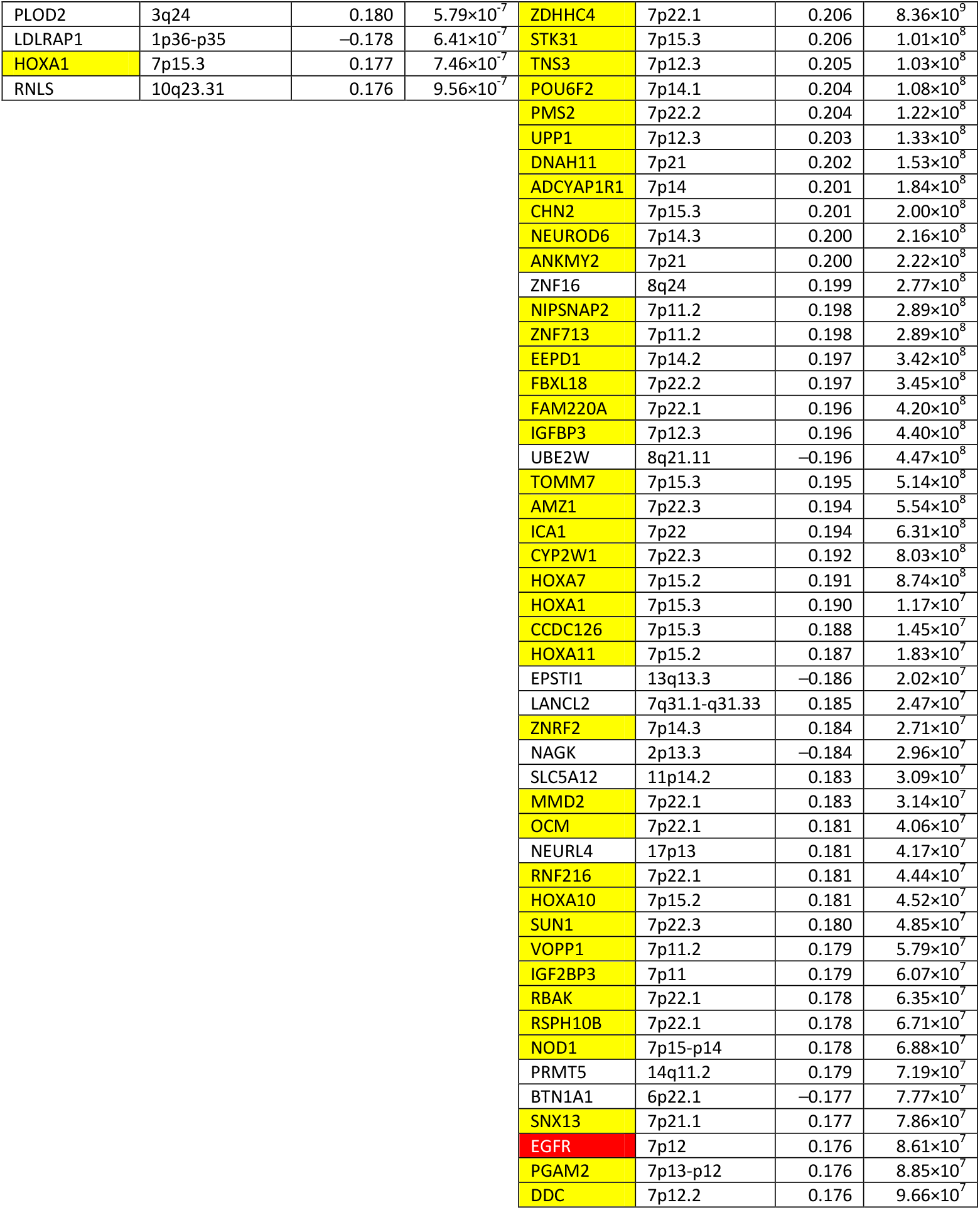
List of genes whose extinction pattern is correlated with those of *AGR2* and *AGR3* (*p* value ranking, down to < 10^−6^) Genes located on 7p are spotted in yellow, oncogenes are in red and TSG are in blue.

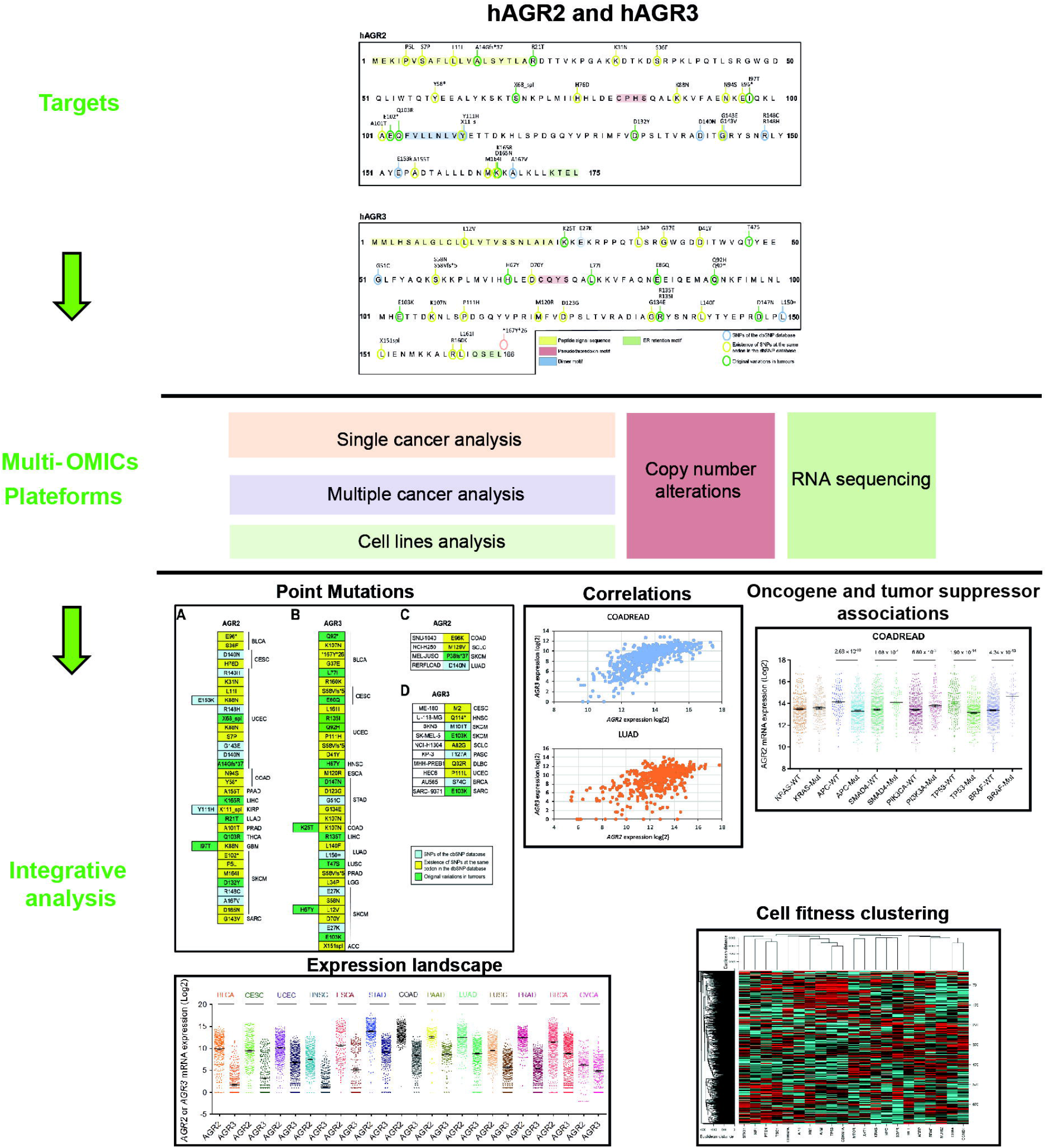

